# A multi-omics approach to identify deleterious mutations in plants

**DOI:** 10.1101/2024.08.22.609273

**Authors:** Omer Baruch, Avraham A. Levy, Fabrizio Mafessoni

## Abstract

Crops lose genetic variation due to strong founder effects during domestication, accumulating and potentially exposing recessive deleterious alleles. Therefore, identifying those deleterious variants in domesticated varieties and their functional orthologs in wild relatives is key for plant breeding, food security and in rescuing the biodiversity of cultivated crops. We explored a machine learning strategy to estimate the impact of new and existing mutations in plant genomes, leveraging multi-omics data, encompassing genomic, epigenomic and transcriptomic information. Specifically, we applied a support-vector-machine framework, previously applied to animal datasets, to published omics data of two important crops of the genus Solanum - tomato and potato - and for the model plant *Arabidopsis thaliana*. We show that our approach provides biologically plausible inferences on the role of mutations occurring in different genomic regions and predictions that correlate with natural genetic variation for the three species, supporting the validity of our estimates. Finally, we show that our estimates outperform existing methods relying exclusively on phylogenetic conservation and not leveraging the availability of omics data for crop species. This approach provides a simple score for researchers to prioritize variants for gene editing and breeding purposes.

## Introduction

Domesticated species tend to have a higher proportion of genetic variation considered to be damaging to fitness^1^. This occurs because, generally, a selected number of individuals, representing a small part of the ancestral population, contribute to the gene pool of the domesticated population, resulting in strong population bottlenecks. In addition, the population undergoing domestication is subjected to strong human-driven selection for specific desirable phenotypes, often through inbreeding, significantly reducing the genetic variation across a range of alleles and often in wide genomic regions surrounding the selected variants because of hitchhiking. The overall reduction in effective population size results in the accumulation of deleterious alleles, potentially even reaching fixation in domesticated cultivars because of hitchhiking along selected variants or because of drift^1–3^. Hence, a primary effort in breeding and crop improvement is that of identifying regions rich in deleterious variants, and alleles carrying potentially advantageous variants in wild relatives, which could potentially be exploited to reintroduce desirable traits in cultivated crops.

Several strategies have been devised to achieve this goal. A common intuition among many of the methods to identify regions under selection is that new mutations in genomic regions of strong importance for the fitness of an individual will result in stronger effects on the phenotypes. As most random mutations are thought to be detrimental, this will result in a higher fraction of new mutations to be purged by natural selection - specifically negative or purifying selection. Hence, these regions will appear more evolutionary conserved across species. Several methods exploit this intuition, which we define as “phylogenetic conservation”. One of the most influential is GERP and its successor GERP++^4^, which quantifies the severity of mutations in terms of Rejected Substitutions, namely, the proportions of mutations that have been rejected by natural selection compared to a neutral expectation based on the phylogenetic relationships between the species analyzed. Another method using this intuition is PhastCons^5^, which employs a Hidden Markov Model - a probabilistic algorithm which scans the genome and assigns different regions to different states, in this case “conserved” or “non-conserved” regions. Another very popular algorithm is PhyloP^5^, a probabilistic method that uses the same intuition as the previous ones and became widely used in literature. Using this method and leveraging multiple whole-genome alignments of hundreds of animal species, recently a large consortium (Zoonomia Consortium) generated single-base resolution characterization of the fitness effects of mutations in mammals, including humans, with implications for conservation biology and clinical studies. A similar effort was focused specifically on primates^6^. Note that while constrained genomic regions usually show a slower rate of evolution and a reduced accumulation of mutations between species, these regions might be under different degrees of constraint over the course of evolution and across different lineages. In animal genomes, it has been found that a substantial fraction of the genome undergoes turnover of selective constraints^5^. Remarkably, no such estimates exist for plants, where possibly, the rate of turnover might be higher due to the high plasticity of plant genomes. Despite this, phylogenetic conservation provides a powerful and promising feature to exploit also for plant studies. Recently, a pipeline to generate multiple whole genome alignments and facilitate the construction of deleteriousness scores based on phylogenetic conservation was developed for plants^7^.

Using a similar intuition to that of phylogenetic conservation, one can in principle use allele frequencies in a population to infer regions under negative selection. Deleterious mutations will in fact usually persist at lower allele frequencies in a population, since natural selection will remove individuals with lower fitness. Hence, different methods used allele frequencies from individual populations - often in conjunction to between-species substitutions - to identify functional regions where mutations are more likely deleterious. One of these approaches - INSIGHT^8^ - has also been applied to plants, developing a tool called greenINSIGHT^9^. This method employs a probabilistic algorithm, pooling all genomic regions sharing a specific feature (e.g. highly methylated regions) and aggregating them to estimate their overall putative deleteriousness. Such a method has been applied to Arabidopsis and rice^9^.

More recently, attempts have been developed to employ machine learning methods aiming at identifying genomic regions under constraints. A recent attempt is that of Benegas^10^, who deployed a method based on Natural Language Processing - thereby redefined as DNA language processing - to find recurrent regions in a genome and closely related ones, with specific features which appear recurrent. Hence, regions are classified as “functional/putatively deleterious” if highly predictable, indicating that a specific sequence does not occur at random, and thus, it is likely, not random. It is important to notice that this pioneering approach is still in its infancy and has to face several limitations: first, especially in plant genomes, highly predictable regions are repeats, which abound in plant genomes; second, so far, this method could only be scaled to genic regions.

Another biological intuition, which has been tackled through the usage of the machine-learning approach, is that for many species, plenty of data is available, including transcriptomic, proteomic and epigenomic data. In principle, such data may provide precious information. For example, it is immediate to think that highly transcribed alleles might be more important for the fitness of an organism than un-transcribed alleles; or that mutations affecting specific protein domains are more important than others. Such information can, in principle, be used to train machine learning algorithms to distinguish between deleterious and neutral mutations. However, we must first define the initial sets of deleterious and neutral mutations to train our algorithm. Since these are usually not known, Kircher et al.^6^ suggested using population genomics principles to build datasets enriched in such types of mutations. Specifically, since mutations that arose and spread to an entire population are unlikely to be deleterious, or at least certainly not lethal and most likely not highly deleterious, such mutations can be used as “proxy-neutral” mutations, i.e. mutations among which the majority is neutral or quasi-neutral. On the other hand, novel mutations have not yet experienced natural selection, and can therefore lead to negative fitness effects. Hence, one could, in principle, build a set of “proxy-deleterious” mutations, i.e. a set enriched in mutations causing negative effects, such non-sense mutations leading to the truncation of proteins necessary for an organism’s survival, by simulating new mutations arising following de novo mutation rates estimated from the genome. Note that these sets do not contain exclusively neutral or exclusively deleterious mutations, but are specifically enriched in these two types of mutations. Using these sets, Kircher et al.^6^ applied a machine learning algorithm – Support Vector Machines^11^ – to learn the genomic, epigenomic and transcriptomic features that distinguish deleterious and neutral mutations in humans (Figure 1). Such an approach has been since expanded and applied to other animal species^12,13^.

Here, we adapted the multi-omics approach developed for humans^6^, to generate maps of fitness effects for three domesticated plants: *Solanum lycopersicum* (tomato), *Solanum tuberosum* (potato) and *Arabidopsis thaliana.* We show that similarly to what is observed in animal genomes, this approach estimates a high contribution of mutations in coding regions to the fitness of organisms compared to intergenic and putatively regulatory ones. We validated this approach by examining the allele frequency of genetic variants in natural populations, showing that our method outperforms those based exclusively on phylogenetic conservation, displaying a higher predictive power of allele frequencies. We tested different approaches - including Support Vector Machines and Artificial Neural Networks - and we report a blueprint to extend this approach to other plant species. Finally, we provide lists of the most deleterious variants that occurred during the domestication of tomato and potato, identifying specific pathways affected during this process.

## Results

### Construction of datasets and predictions

We first focused on building maps of fitness effects for tomato. We built an alignment using 59 species, with 36 representatives of Solanaceae, out of which 28 representatives of the genus Solanum (Supplementary Table 1), using the *msa* pipeline^7^. For tomato, we used the genome Heinz SL5.0^14^. Using the same pipeline, we computed conservation scores, specifically GERP++^4^, PhyloP^5^ and PhastCons^5^. For PhyloP, we computed conservation scores without including the reference SL5.0^14^ and genomes that putatively belonged to the species *S. lycopersicum* or provided genetic introgression material to *Solanum lycopersicum* (*S. lycopersicum var. cerasiforme* and *S. pimpinellifolium*), following the procedure described by Kircher et al.^15^, and after an initial evaluation in which we assessed that including the reference was biasing inferences (see Appendix). We then computed other genomic features (e.g. GC content) and features describing amino-acid substitutions using SIFT and various substitution matrices (BLOSUM62, grantham). We then collected transcriptomic and epigenomic features. Specifically, we obtained histone modification, methylation, short RNAs, DNA hypersensitivity and chromatin accessibility data available for *S. lycopersicum*. Data that were collected and mapped for different genome assemblies other than SL5.0 (e.g. SL2.5) were processed *de novo* and mapped to SL5.0, to avoid artifacts and missing data that might arise when lifting tracts from one assembly to another. Details of these steps are detailed in the Methods section. Features were normalized as described in the Methods. A flowchart of the procedure is shown in Figure 2.

**Figure 1.**
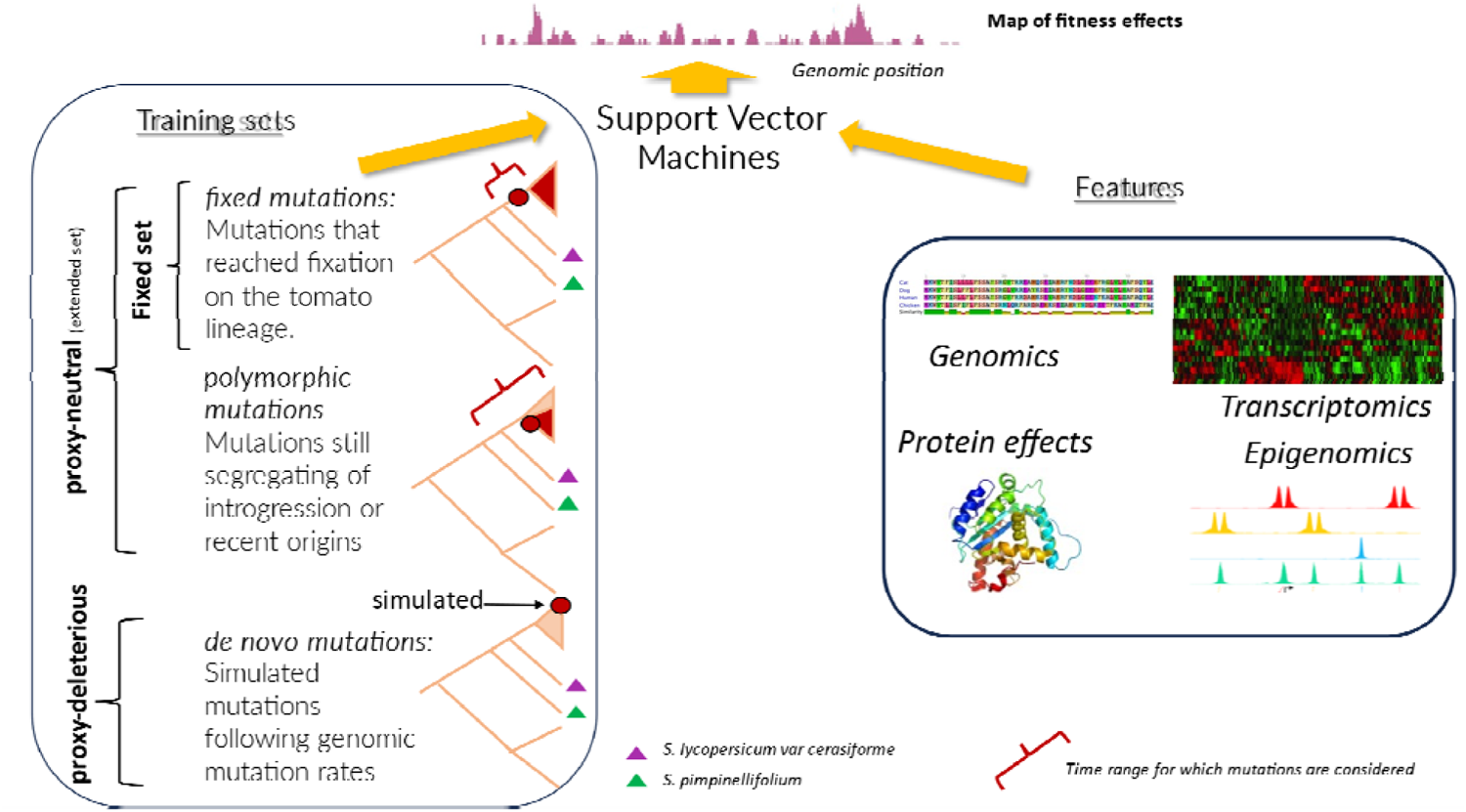
General overview of the approach. A training set (left panel) is built with mutations evolved in the tomato lineage, with both fixed and polymorphic positions at which the derived allele is found in *S. lycopersicum* with respect to its ancestor with *S. pimpinellifolium* (considered as proxy-neutral, i.e. mutations enriched in non-deleterious mutations). This allows also for mutations introgressed from *S. pimpinellifolium* in the *S. lycopersicum var. lycopersicum* (but differing from *S. lycopersicum var. cerasiforme*). Proxy-deleterious mutations, i.e. mutations enriched in deleterious mutations, are built via simulations mimicking *de novo* mutation rates in tomato. The same procedure is applied for *A. thaliana* and potato. Fixed mutations are used for additional trainings, of which validation plots are in Supplementary Figure 1 for tomato and Supplementary Figure 2 for *Arabidopsis*. In the Training Sets panel (left), the phylogenetic relationships between species are represented in pink, with the time and phylogenetic placement of an example-derived mutation included in that training set as a red dot and the descending lineages carrying the same mutation in the same color. Variation within the focal species (here tomato) is shown as a triangle, colored in red proportionally to the amount of individual carrying the derived mutation. Features informative of different genomic regions and mutations (right panel) are collected and used to train a Support Vector Machine which learns to discriminate between the two training sets, by learning which features are “typical” of neutral and deleterious mutations.

We then generated the training sets, based on the Heinz SL5.0^14^ reference genome, creating mutations that we assigned to two groups: proxy-neutral and proxy-deleterious mutations. Proxy-neutral mutations were chosen as mutations that evolved on the tomato lineage (Figure 1) in respect to the ancestral state, here defined as the allele shared by the majority of genomes in the whole-genome alignment under the condition that either *S. lycopersicum var. cerasiforme* a wild ancestor of the domesticated tomato *S. lycopersicum var. lycopersicum* or *S. pimpinellifolium* shared this allele – to avoid fixation events that occurred at deeper phylogenetic scales. Positions for which the ancestral state could not be inferred unambiguously using this criterion were excluded from the analyses. Note that this set includes both mutations that are polymorphic and that reached fixation in the *S. lycopersicum* lineage. We also analyzed fixed mutations alone, and to do this we excluded alleles observed as polymorphic in the population data^14^. Proxy-deleterious mutations were generated by simulating a set of new mutations mimicking as close as possible *de novo* mutations, by keeping the spectrum of mutations ( the relative proportions of the different types of nucleotidic substitutions, e.g. A to C) and local mutation rates the same as to those observed in respect to the common ancestor with *S. pimpinellifolium.* We achieve this through a permutation-based scheme, different from that employed by Kircher et al., and detailed in the Methods section.

**Figure 2.**
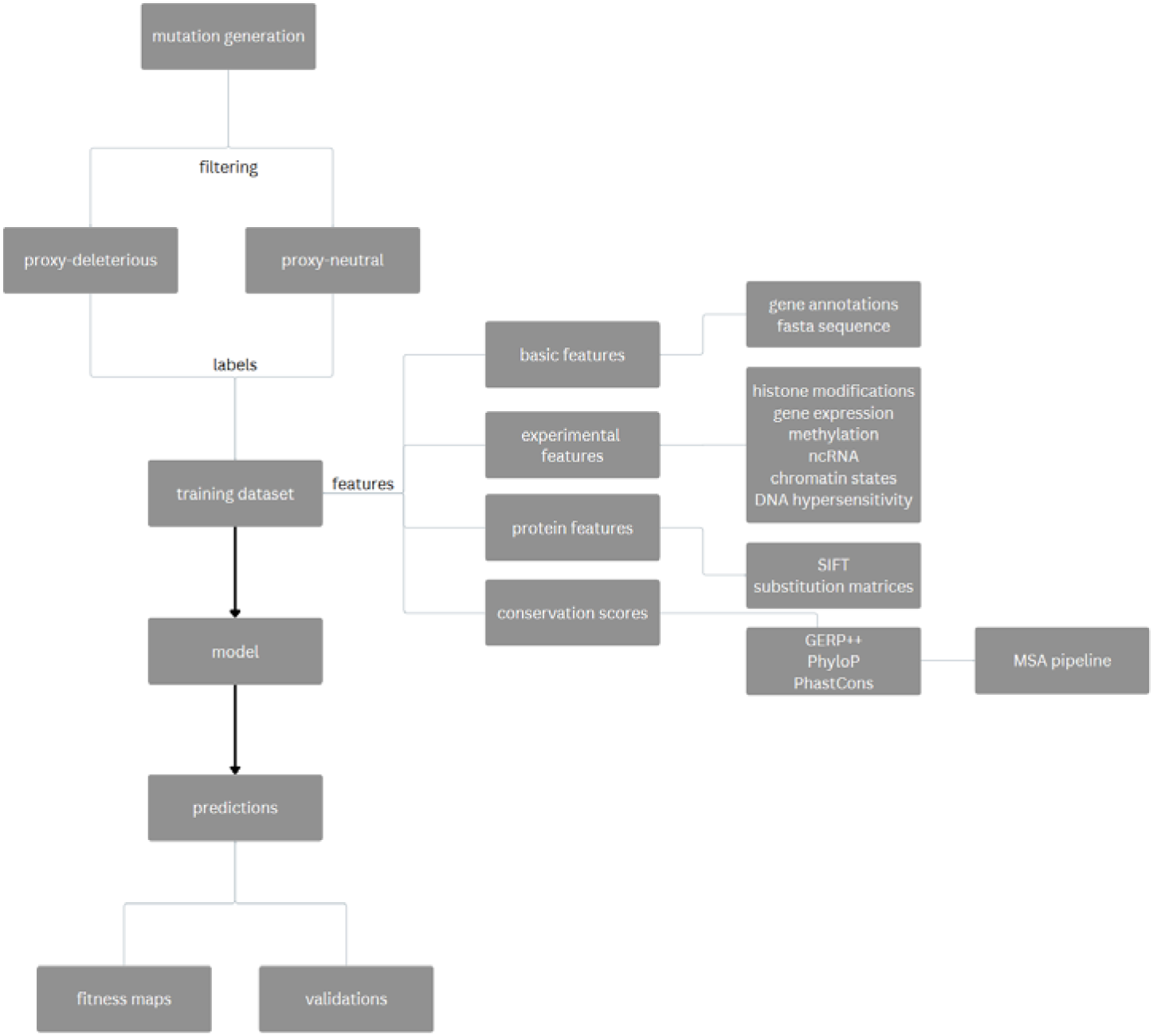
A flowchart describing the procedure of constructing our dataset.

We then applied the same approach to two other species: *A. thaliana,* a model plant for which there is plenty of data available and thus provides an optimal benchmark for this approach, and potato (*S. tuberosum). S. tuberosum* is another important crop in the Solanum genus, and also provides a test case for plants for which less data (both epigenomic and in terms of genetic variation) is available.

For *A. thaliana* we used the assembly TAIR10^16^ and we built a whole genome multiple phylogenetic alignment and conservation scores using the *msa* pipeline^7^ as for tomato. The species used for the alignment are shown in Supplementary Table 2. The annotation and epigenomic data collected are reported in the Methods section. The training set was built using the same criterion as tomato: proxy-neutral mutations were selected as mutations that went to fixation in the focal lineage (here *A. thaliana*) with respect to the ancestral state, here defined a the allele shared by the majority of genomes in the whole-genome alignment under the condition that either *A. suecica* or *A. lyrata* shared this allele – to avoid fixation events that occurred at deeper phylogenetic scales. Positions for which the ancestral state could not be inferred unambiguously using this criterion were excluded from the analyses. As with *S. lycopersicum,* we used a proxy-neutral mutation set. We also included preliminary validation plots for a model trained on a fixed neutral set (Supplementary Figure 2). Proxy-deleterious mutations were also generated using the same scheme as tomato, by the positions of alleles that differed between *A. thaliana* and *A. arenosa*.

For potato, we used the assembly DM6.1^17^, and a multiple whole-genome alignment including the species reported in Supplementary Table 3. The annotation and epigenomic data collected are reported in the Methods section. The training set was built using the same criterion as described above: proxy-neutral mutations were selected as mutations that went to fixation in the focal lineage (here *S. tuberosum*) with respect to the ancestral state, here defined as the allele shared by the majority of genomes in the whole-genome alignment under the condition that either *S. stenototum* or *S. verrucosum* shared this allele - to avoid fixation events that occurred at deeper phylogenetic scales. Positions for which the ancestral state could not be inferred unambiguously using this criterion were excluded from the analyses. Proxy-deleterious mutations were also generated using the same scheme as tomato, by the positions of alleles that different between DM6.1 and *S. okadae*, taken as an outgroup for the *S. tuberosum* clade. In an effort to make a fixed neutral set, we obtained data from the *solomics* datasets^18^. This dataset contains 432 accessions, mostly of wild-potato genomes. We retained accessions defined as *S. goniocalix, S. stenototum and S. phureja.* However, after lifting the limited dataset (47k positions) we were left with an extremely limited set (11k positions). Thus the potato model is effectively trained using a proxy-neutral set as described.

To test that our training sets were truly enriched in putatively-neutral mutations we examined the proportion of non-synonymous versus synonymous changes in protein coding regions. Specifically, if our proxy-neutral training set was not enriched in neutral mutations we would observe the same proportion of mutations in the first, second and third codon positions. First, we examined these proportions when population data was used to retain only fixed mutations as described above. We observe a significantly higher proportion (p-value<10^∧^16 for chi square test) of mutations in the third codon (Table 1). The highest ratio is observed for Arabidopsis, while the lowest for tomato. Thus, we conclude that our fixed-neutral training sets are enriched for neutral mutations as desired. We also computed the same quantity without excluding mutations that were observed as polymorphic in population data. Unexpectedly, the fraction of proxy-neutral mutations in the third codon was higher than in the previous sets for tomato and Arabidopsis, which have the largest amount of polymorphic data. This can be potentially explained by the fact that extensive introgression exists between our focal taxa and the sister species. Hence, removing polymorphic variants will preferentially exclude such introgression events, which might bias our datasets, particularly if introgression occurs more frequently in neutral regions - which is often observed in natural populations. Hence, for subsequent analyses, we used data in which polymorphic positions in population data were included (described as the proxy-neutral set). Note that this is particularly useful for potential applications in species for which extensive population data is not available. In addition, excluding polymorphic positions would reduce the size of the proxy-neutral training datasets, by ∼10% and 20% in tomato and Arabidopsis, respectively. Note however, that we see a larger enrichment in intergenic regions when excluding polymorphisms for *S. lycopersicum* (Supplementary Table 4).

**Table 1.** The proportion of mutation in the proxy-neutral sets found in the different codon-positions (Codon position). Mutations in the 3rd codon are expected to be enriched in synonymous changes. Proxy-neutral mutations were filtered to exclude polymorphic mutations in population data (Fraction fixed in population data) or not (Fraction).

| Species | Codon position | Fraction proxy-neutral | Fraction fixed-neutral |
| --- | --- | --- | --- |
| *Arabidopsis thaliana* | 1 | 0.2790 | 0.2817 |
|  | 2 | 0.2523 | 0.2533 |
|  | 3 | 0.4686 | 0.4649 |
| *Solanum lycopersicum*<br>(tomato) | 1 | 0.3171 | 0.3324 |
|  | 2 | 0.3025 | 0.2982 |
|  | 3 | 0.3803 | 0.3692 |
| *Solanum tuberosum*<br>(potato) | 1 | 0.2975 | - |
|  | 2 | 0.2830 | - |
|  | 3 | 0.4193 | - |

We trained both a Deep Learning model^19^ and a Support Vector Machine model^11^ (SVM) to discriminate between proxy-neutral and proxy-deleterious mutations. A comparison of the performance of the two machine learning approaches (Supplementary Table 5) led us to opt for SVMs. The SVM model outputs scores between 0 and 1, which we define as “raw scores”, where 1 indicates more deleterious mutations. To allow for a better visualization of highly deleterious variants we also apply a PHRED-like scaling of the scores:

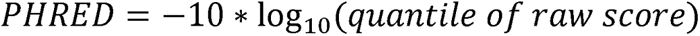

so that a score of 10 indicates that a mutation has an effect corresponding to the top 10% of all novel mutations, a score of 20 to the top 1%, a score of 30 to the top 10^-3^ and so on.

In Figure 3 we show the proportion of mutations of different types stratified by the raw deleteriousness score obtained from the SVM model for 4.41M novel mutations simulated based on the genomic distribution of proxy-deleterious mutations on the *S. lycopersicum* lineage. In Figure 3a we show the distribution of mutations in genomic regions along the quantiles of our prediction scores. The plot shows the relative abundance of each genomic region as we normalize each mutation type to have the same overall abundance. We observe how predictions correspond to biological expectations: mutations in coding regions appear among the most deleterious, with protein changing mutations being overwhelmingly more deleterious than synonymous ones, and mutations resulting in the gain or loss of stop codons being even more abundant among the most deleterious mutations. As expected, intergenic and intronic regions have in general much smaller effects than coding mutations (Fig 3b), with the vast majority of intergenic regions having PHRED-scaled scores smaller than 20, i.e. their effect is smaller than the top 1% deleterious mutations, though because of their sheer amount, some intergenic regions are predicted to have strong effects on fitness (PHRED-scaled scores larger than 30, i.e., their effect is stronger than the top 0.1% most deleterious variants) emphasizing the advantage of a genome-wide approach. Similarly to intergenic mutations, some synonymous mutations appear to have strong effects. Interestingly, we do see some synonymous mutations in the top deleterious variants, suggesting these mutations have a strong effect on fitness, as will be discussed later.

For *Arabidopsis* and potato, we observe similar patterns to those observed for tomato. Intergenic and synonymous mutations appear as the most neutral mutations in all species, while non-synonymous mutations have stronger effects than either, and mutations affecting stop or start codons have the most severe effects. Note that due to the smaller number of potential mutations resulting in stop-codon gains or start-codon losses, particularly for the latter, and their strong effects, they become much more visible in the plots showing PHRED-scaled scores, which zoom on highly deleterious mutations (Figures 3b, 4b and 5b).

We also detect some differences between species. Notably, both *Arabidopsis* and potato show a stronger distinction between non-synonymous mutations and the more deleterious mutations resulting in gains of stop codons. *Arabidopsis* also shows proportionally a larger number of coding mutations compared to intergenic, which is mostly visible in the larger number of neutral synonymous variants. We hypothesized that these differences are likely due to differences in the overall composition of the genomes, either because of the actual genome composition or become of our alignments. For example, the analyzed *Arabidopsis* genomic dataset is much richer in genes than either tomato or potato (∼50%, 20% and 27% respectively). In addition, the training sets size for *Arabidopsis* and potato were larger than that of tomato when adjusted to their genome size (∼4%, 2% and 0.5% respectively). These two combined effects potentially resulted in a higher resolution in estimating the effects of low abundance mutation classes, like mutation affecting stop codons.

**Figure 3.**
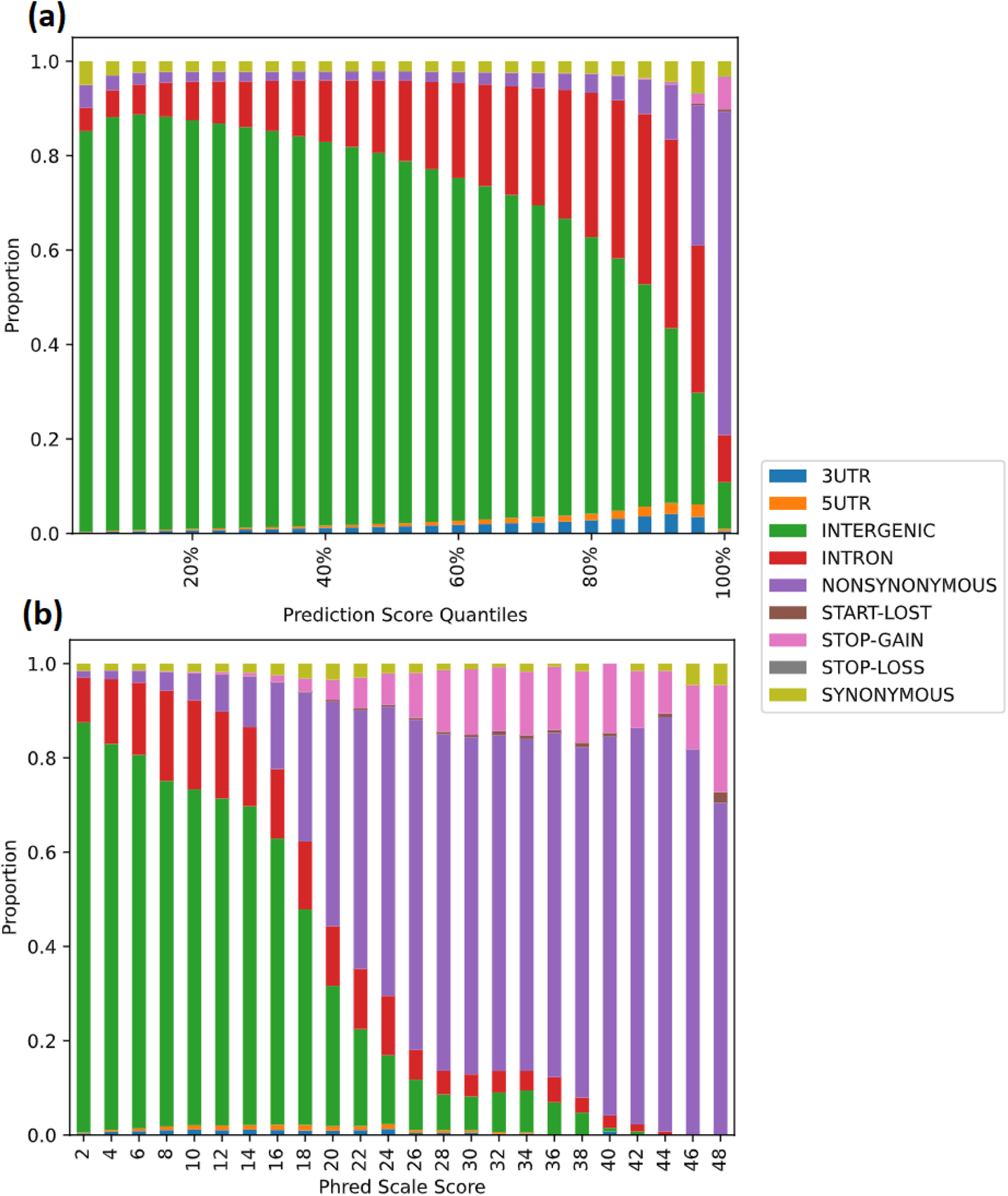
Distribution of the effects of 4.41M proxy-neutral mutations for *S. lycopersicum*. Different colors correspond to different types of genomic regions according to the annotations reported for SL5.0 as described in the legend. The prediction scores were divided into percentiles and plotted in 25 bins spanning 4 percentiles (4%), with increasing deleteriousness. (a) The proportion of different genomic regions (y-axis) in each score quantile (x-axis) of the raw-score predicted by the SVM model is shown for increasingly more deleterious variants (larger quantiles). (b) The proportion of different genomic regions (y-axis) for different PHRED-scaled scores is shown for increasingly more deleterious scores. PHRED-scaled scores are grouped in bins of 2 (x-axis).

**Figure 4.**
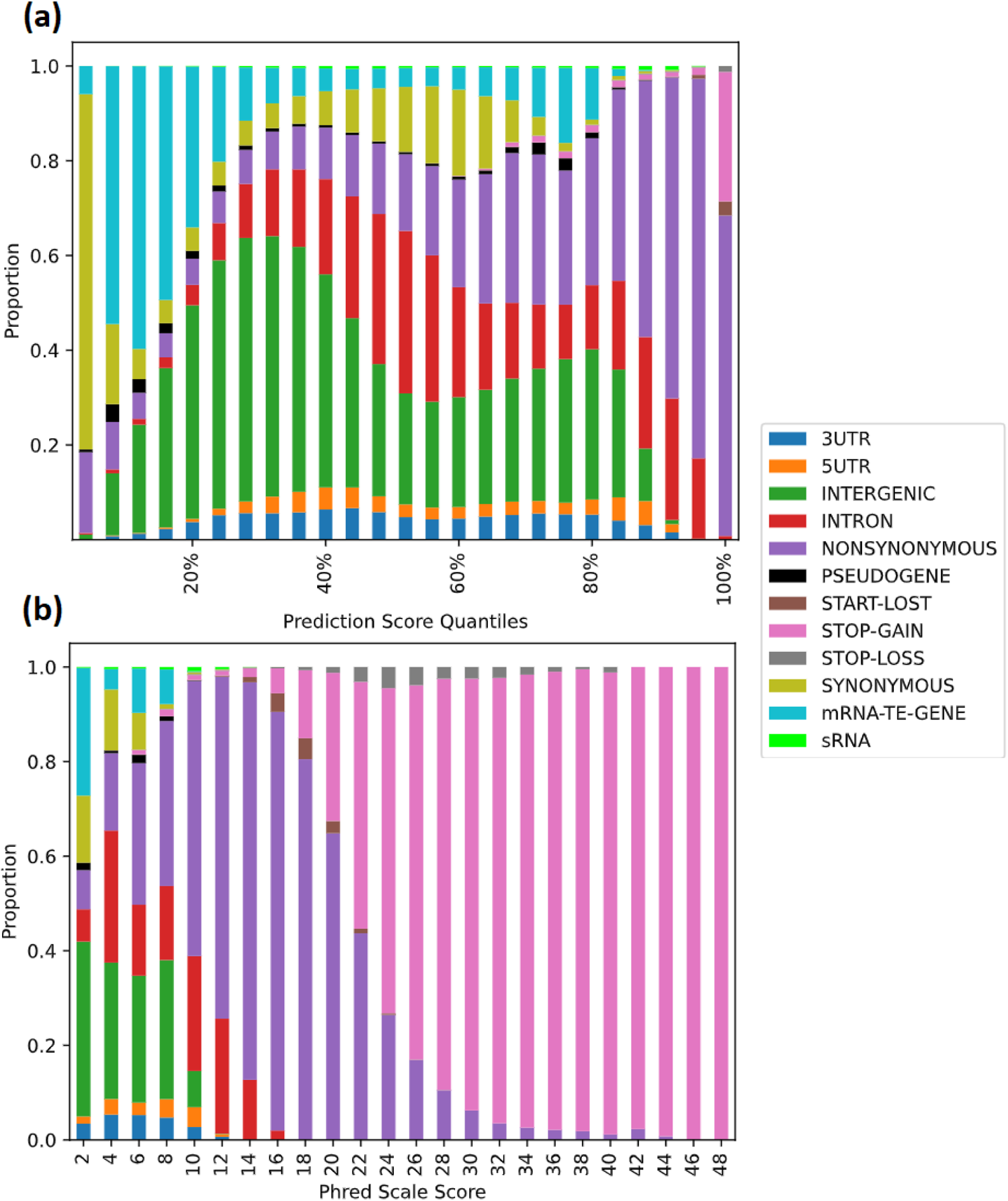
Distribution of the effects of 5.38M new mutations by annotation for *A. thaliana*. Different colors correspond to different types of genomic regions according to the annotations reported for TAIR10 as described in the legend. The prediction scores were divided into percentiles and plotted in 25 bins spanning 4 percentiles (4%), with increasing deleteriousness. (a) The proportion of different genomic regions (y-axis) in each score quantile (x-axis) of the raw-score predicted by the SVM model is shown for increasingly more deleterious variants (larger quantiles). (b) The proportion of different genomic regions (y-axis) for different PHRED-scaled scores is shown for increasingly more deleterious scores. PHRED-scaled scores are grouped in bins of 2 (x-axis).

**Figure 5.**
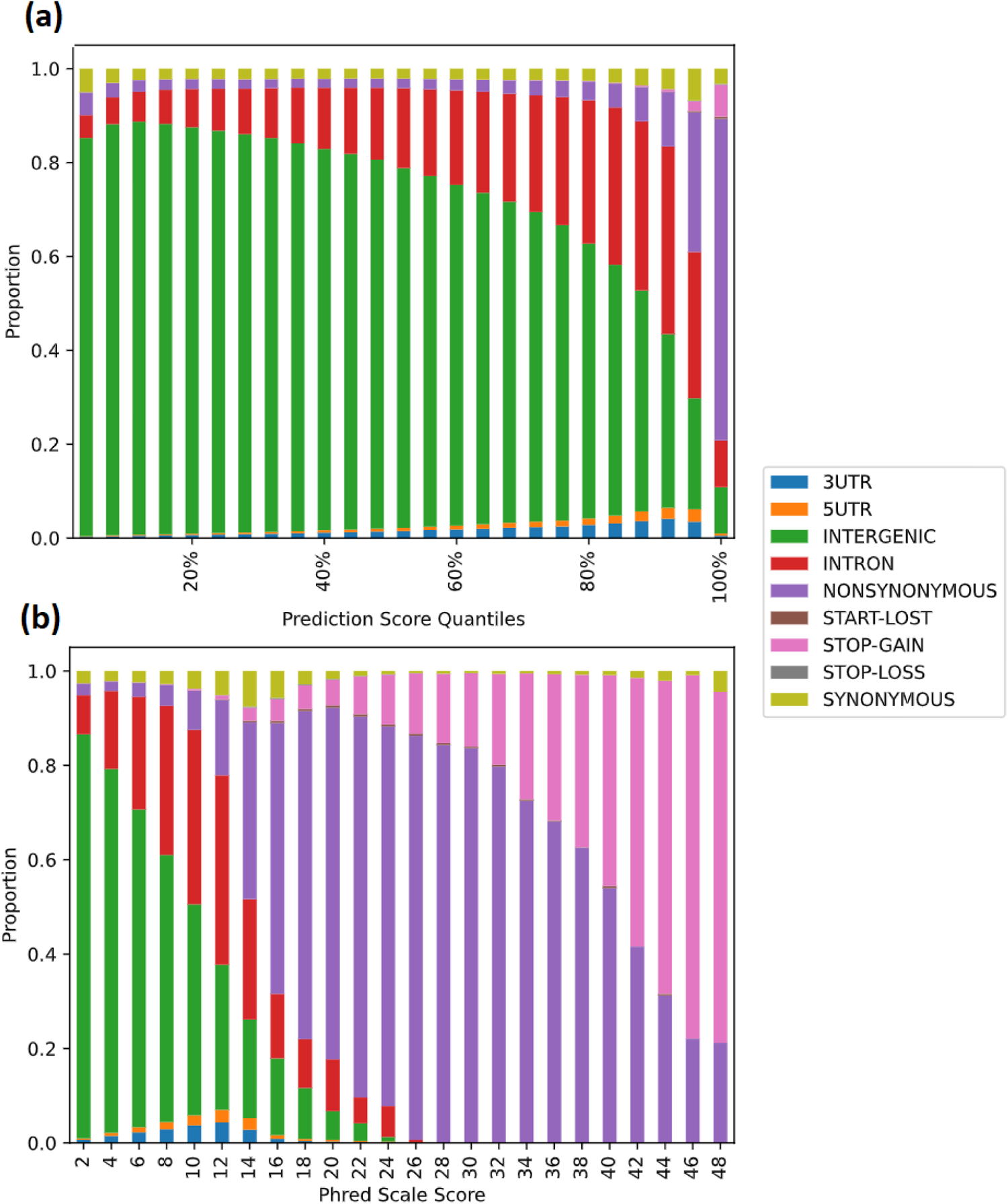
Distribution of the effects of 11.30M proxy-deleterious mutations for *S. tuberosum*. Different colors correspond to different types of genomic regions according to the annotations reported for DM 6.1 as described in the legend. The prediction scores were divided into percentiles and plotted in 25 bins spanning 4 percentiles (4%), with increasing deleteriousness. (a) The proportion of different genomic regions (y-axis) in each score quantile (x-axis) of the raw-score predicted by the SVM model is shown for increasingly more deleterious variants (larger quantiles). (b) The proportion of different genomic regions (y-axis) for different PHRED-scaled scores is shown for increasingly more deleterious scores. PHRED-scaled scores are grouped in bins of 2 (x-axis).

### Validation of the method and comparison with other approaches

To validate our method, we tested whether it could predict the frequency of existing mutations in natural populations. In fact, deleterious alleles are expected to segregate at lower allele frequencies than neutral ones. Hence, a negative correlation between measures of deleteriousness per allele and allele frequencies is expected. For tomato, using the Zhou^14^ dataset, our score shows a highly significant negative (p-value < 10^-16^) correlation coefficient with allele frequencies (Figure 6). We thus used this measure to compare our method to other existing ones (Figure 7, Table 2). We observe that our model provides the strongest correlation with derived allele frequencies (-0.161) when compared to other common methods like PhyloP (-0.083) and GERP (-0.063). PhyloP, for example, shows a bimodal pattern, with variants estimated to have an effect between the 40% and 60% percentile of the score having mean allele frequencies lower than those with very high effects. In addition, the most neutral bin (0-10%) have mean allele frequency ∼0.09 while the most severe frequency 0.04. Our predictions show a more gradual pattern and a larger difference in mean allele frequency between bins with the highest allele frequency (0.14 for bin 10-20%) the most severe mutations (mean allele frequency 0.03). Note, however, that the pattern is complex: the lowest quantile bin of the score (0-10% most neutral mutations) shows lower mean allele frequency than higher bins of the score. We hypothesize that this could be due to ancestral misassignment - alleles that are actually at low allele frequencies, but that are assigned as ancestral rather than derived - or due to low-frequency introgression from wild relatives of advantageous alleles lost during domestication, which is pervasive for tomato. We found this pattern to be present also when restricting our analyses to the 350 plants with the lowest divergence from the Heinz reference genome to exclude introgressed accessions and accessions of *Solanum lycopersicum var. cerasiforme*. Consistent with the hypothesis that this pattern could be due to the extensive introgression in tomato and typical of many domesticated species, Arabidopsis shows a smoother and more monotonic pattern. The population data used was from the 1001 Genomes project^20^. Note that GERP shows the highest accuracy, measured as the proportion of rare mutations with higher deleteriousness scores - under the assumption that deleterious mutations are most often rare - while our method shows high accuracy (the highest in tomato) compared to other methods. We did not compute these measures for potato as only 11.7K variants were available in the population data of this species after lifting the population dataset to the DM6.1 assembly, resulting in an unsignificant analysis.

**Table 2.** Performance of different methods to estimate maps of fitness effects in *S. lycopersicum* and *A. thaliana*. Correlation was computed as Spearmans’ rank correlation. Accuracy was computed as the proportion of True Positives (TP) over total classifications, i.e. TP/(TP+FP+TN+FN). Positives (P) were defined as mutations with PHRED-scaled score higher than 30, and negatives as mutations with lower scores. Under the assumption that low-frequency derived alleles are more often deleterious, we defined True Positives as mutations with a PHRED-scaled score higher than 30 and allele frequencies lower than 0.01. Contrarily, False Positives (FP) were defined as mutations with a PHRED-scaled score higher than 30 and allele frequencies and derived allele frequencies higher than 0.1.

|  | Model | Accuracy | Correlation |
| --- | --- | --- | --- |
| *Arabidopsis thaliana* | GERP | 0.619 | -0.110 |
|  | PhyloP | 0.613 | -0.116 |
|  | PHAST | 0.608 | -0.126 |
|  | SVM | 0.617 | -0.138 |
| *Solanum lycopersicum (tomato)* | GERP | 0.803 | -0.063 |
|  | PhyloP | 0.802 | -0.083 |
|  | PHAST | 0.800 | -0.084 |
|  | SVM | 0.810 | -0.161 |

**Figure 6.**
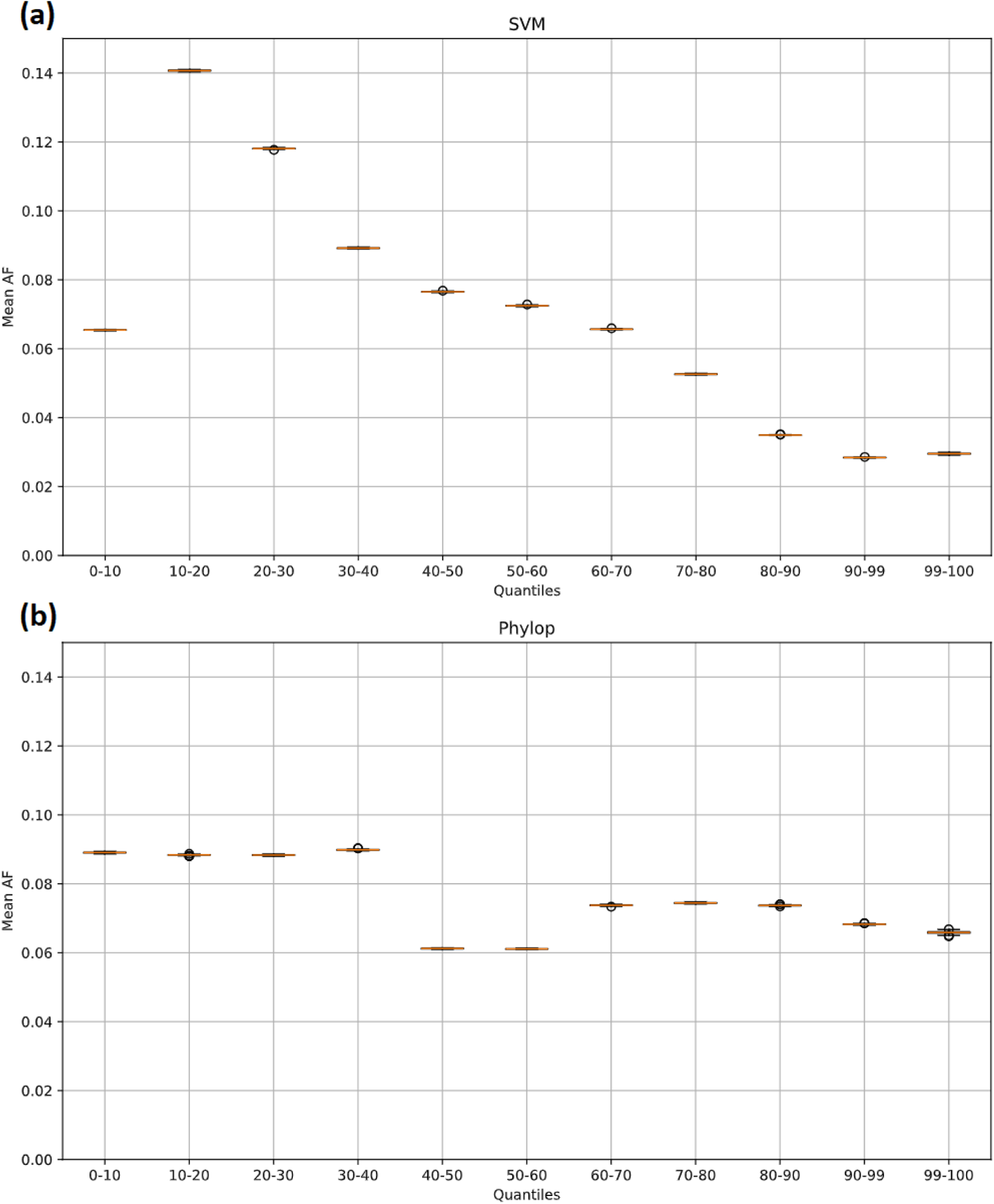
Mean allele frequency (AF, y-axis) for different quantiles of the scores computed by our model (a) and Phylop (b) for tomato. Uncertainty for the mean allele frequency was computed by bootstrapping the alleles in the population data collected by Zhou et al.,2022.

**Figure 7.**
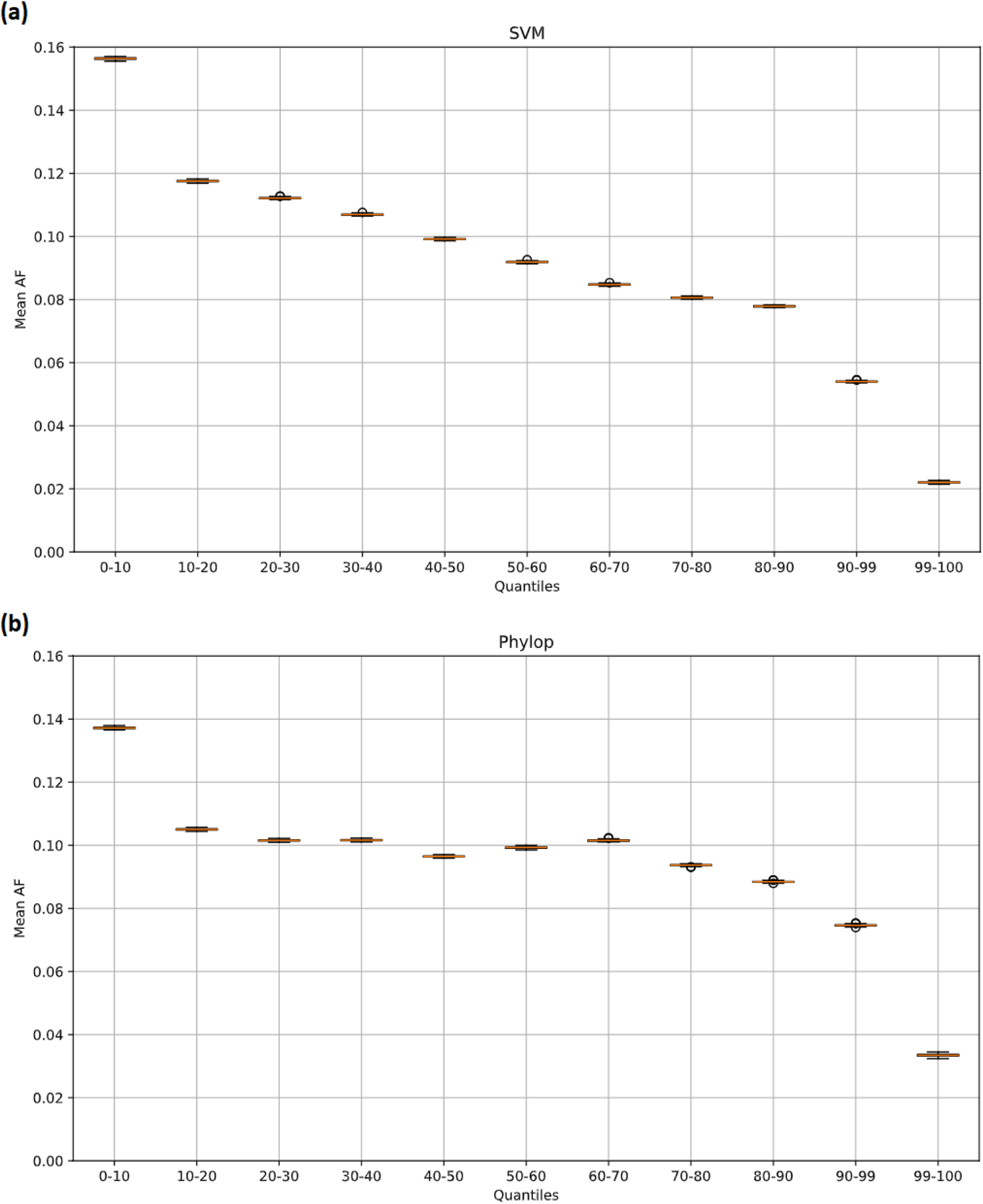
Mean allele frequency (AF, y-axis) for different quantiles of the scores computed by our model (a) and Phylop (b) for *A. thaliana*. Uncertainty for the mean allele frequency was computed by bootstrapping the alleles in the 1001 Genomes dataset.

### Application of the deleteriousness score to investigate the effect of genetic variation in tomato, potato and *Arabidopsis*

We then applied our method to predict the effect of the genetic variants observed in the populations of Arabidopsis and tomatoes. Potatoes were excluded from this analysis due to the small size of the dataset documenting genetic variation for this species (only 11.7k variants, versus the 10.7M for *A. thaliana* and 31.5M for tomato). To do this, we took the top 1000 most deleterious variants and tested whether any pathway or cellular function is enriched among the genes carrying such variants using the program topGO^21^, which has been applied repeatedly to tomato^22,23^. We caution, however, that GO tests performed with topGO are possibly anticonservative, as it does not adjust for biased background gene sets. For variants falling out of genes we considered the gene closest to the genetic variant. In Table 3 and Table 4 we report the top 20 categories for *A. thaliana* and tomato, respectively. For both plants we see the term “catalytic activity” to be overrepresented among polymorphic variants. However, the specific type of pathways involved differ for Arabidopsis and tomato. The former shows a ∼5x enrichment of polymorphic variants in or close to genes involved in heme, iron and ion-binding, together with other genes involved in redox activities. For tomato, we see a general enrichment in nucleotide and ATP metabolism, but also in specialized metabolism, e.g. salicylic acid glucosyltransferase and quercetin 3-O-glucosyltransferase activity.

**Table 3.**
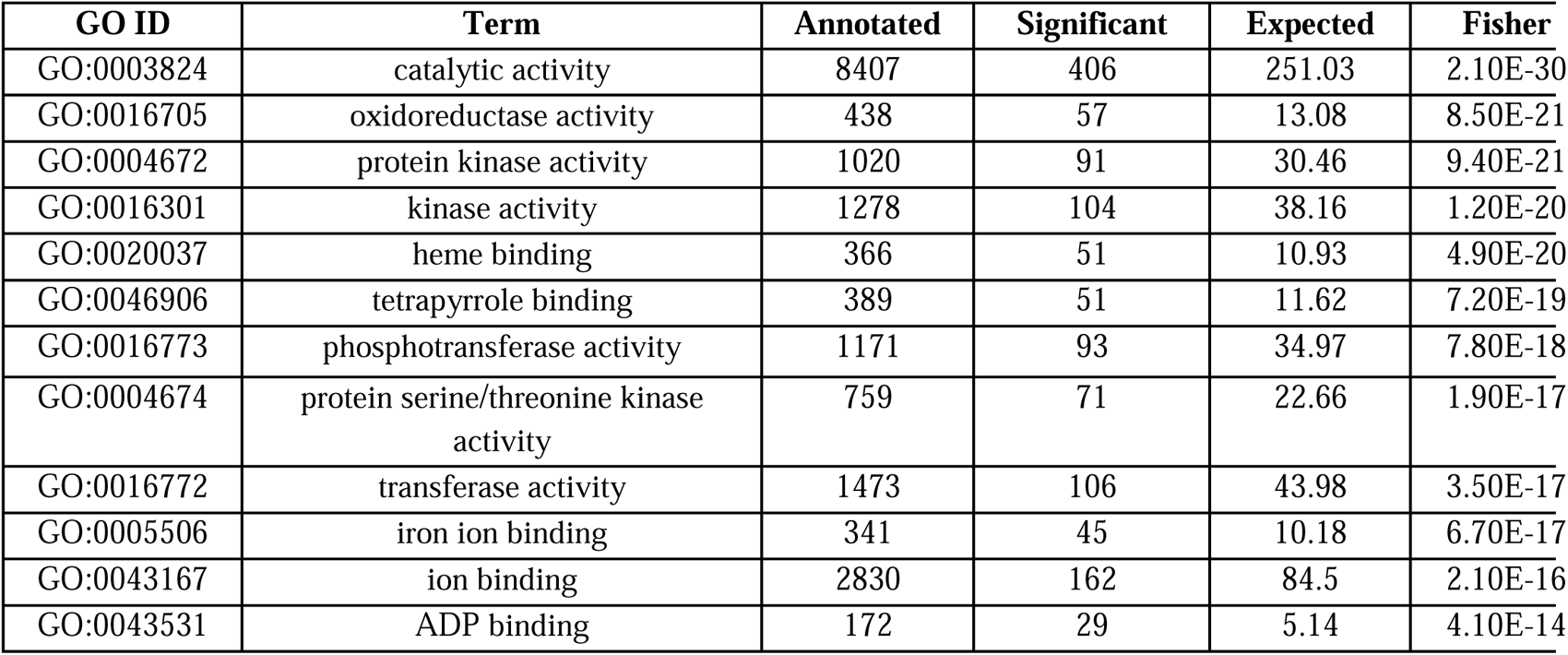
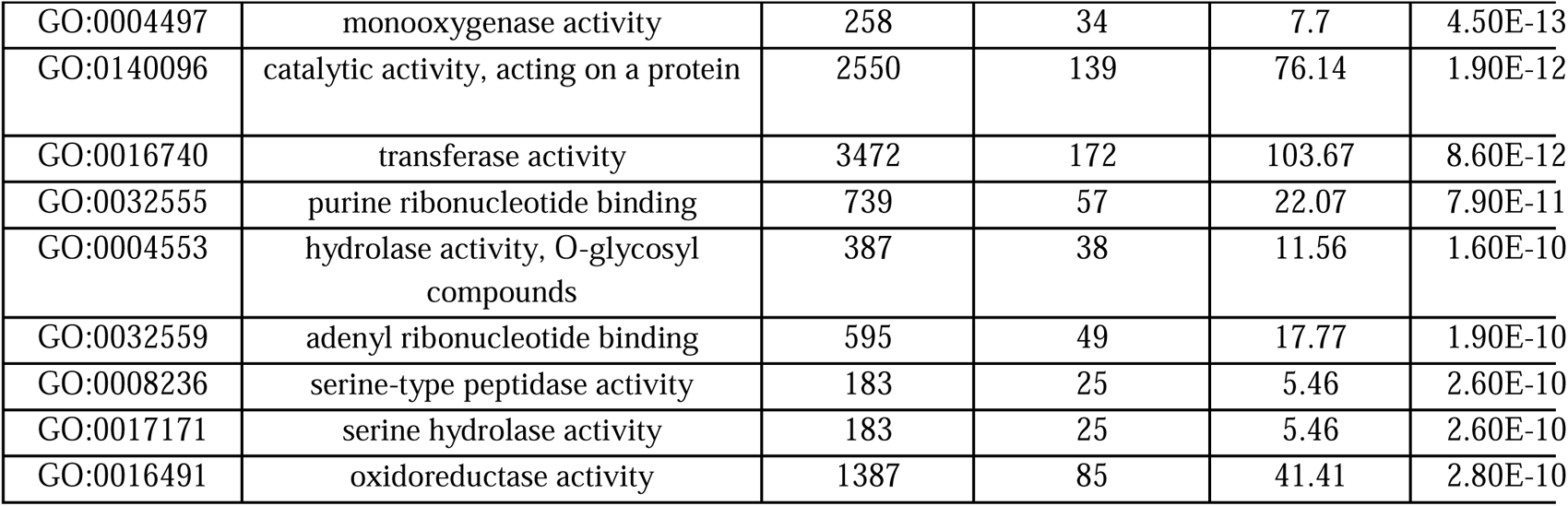
Top 20 gene ontology categories for the genes carrying the most deleterious polymorphic variants in the 1001 Genome dataset of *A. thaliana*. The columns describe the GO ID (GO ID), a brief description of the ontology (Term), the total number of annotated (Annotated), significant (Significant) and expected (Expected) hits for each specific ontology, and the p-value (Fisher). Higher Significant than Expected indicate an enrichment.

**Table 4.**
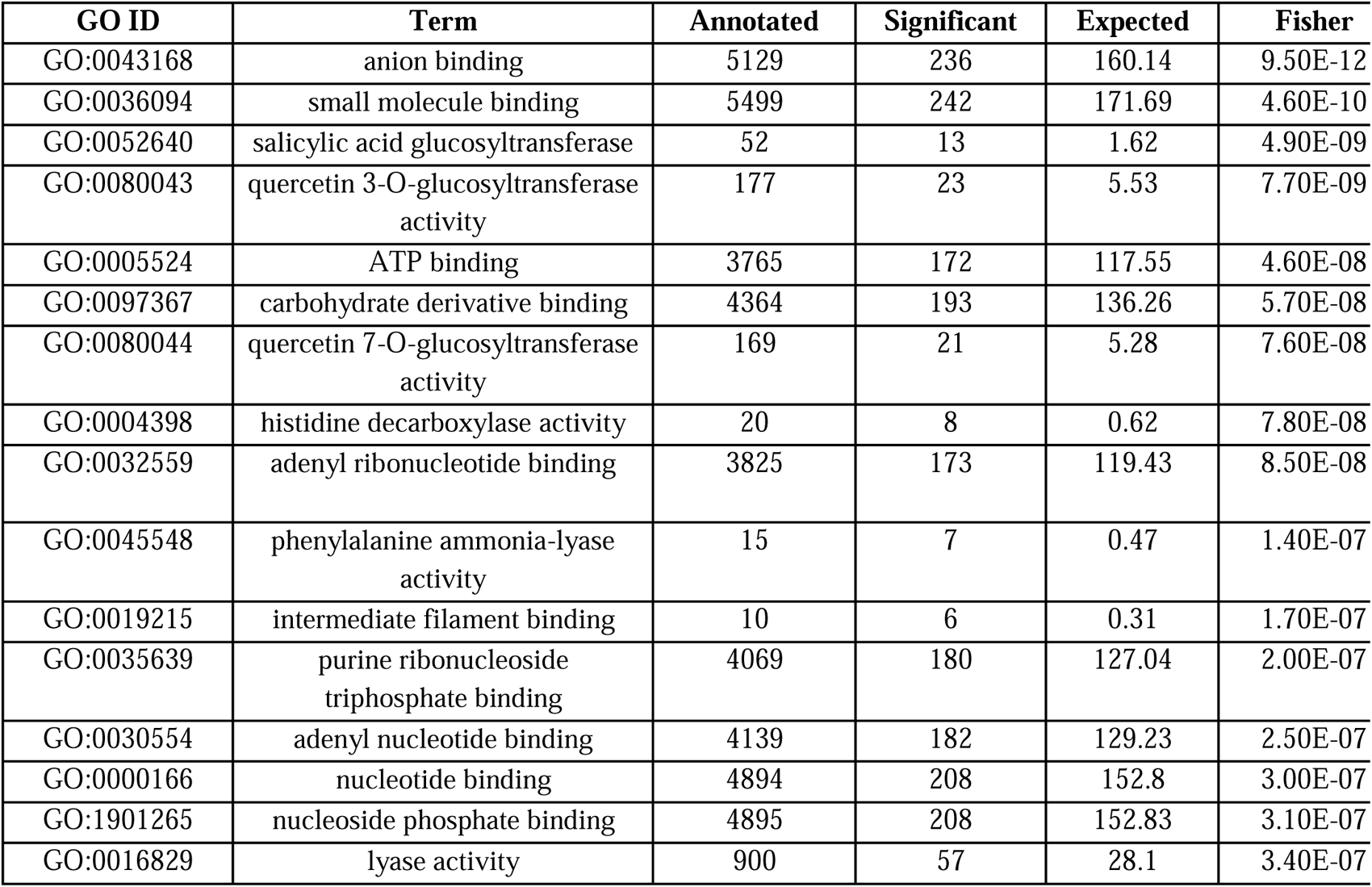
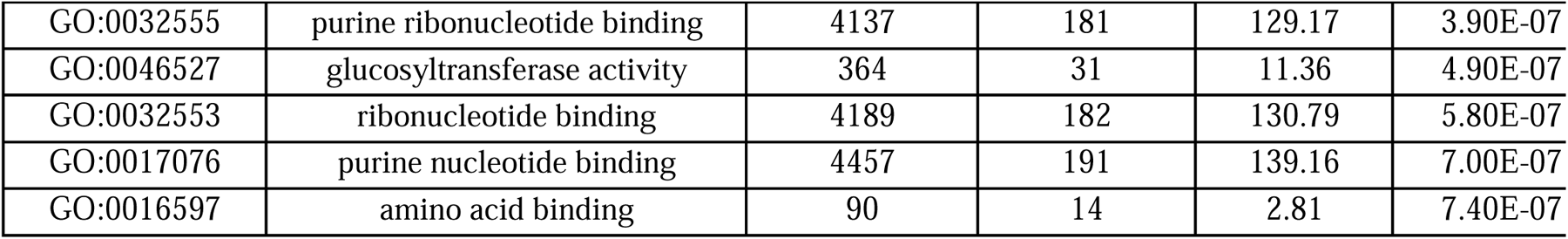
Top 20 gene ontology categories for the genes carrying the most deleterious polymorphic variants in the solomics collection of tomatoes (SL5.0). The columns describe the GO ID (GO ID), a brief description of the ontology (Term), the total number of annotated (Annotated), significant (Significant) and expected (Expected) hits for each specific ontology, and the p-value (Fisher). Higher Significant than Expected indicate an enrichment.

| GO ID | Term | Annotated | Significant | Expected | Fisher |
| --- | --- | --- | --- | --- | --- |
| GO:0043168 | anion binding | 5129 | 236 | 160.14 | 9.50E-12 |
| GO:0036094 | small molecule binding | 5499 | 242 | 171.69 | 4.60E-10 |
| GO:0052640 | salicylic acid glucosyltransferase | 52 | 13 | 1.62 | 4.90E-09 |
| GO:0080043 | quercetin 3-O-glucosyltransferase activity | 177 | 23 | 5.53 | 7.70E-09 |
| GO:0005524 | ATP binding | 3765 | 172 | 117.55 | 4.60E-08 |
| GO:0097367 | carbohydrate derivative binding | 4364 | 193 | 136.26 | 5.70E-08 |
| GO:0080044 | quercetin 7-O-glucosyltransferase activity | 169 | 21 | 5.28 | 7.60E-08 |
| GO:0004398 | histidine decarboxylase activity | 20 | 8 | 0.62 | 7.80E-08 |
| GO:0032559 | adenyl ribonucleotide binding | 3825 | 173 | 119.43 | 8.50E-08 |
| GO:0045548 | phenylalanine ammonia-lyase activity | 15 | 7 | 0.47 | 1.40E-07 |
| GO:0019215 | intermediate filament binding | 10 | 6 | 0.31 | 1.70E-07 |
| GO:0035639 | purine ribonucleoside triphosphate binding | 4069 | 180 | 127.04 | 2.00E-07 |
| GO:0030554 | adenyl nucleotide binding | 4139 | 182 | 129.23 | 2.50E-07 |
| GO:0000166 | nucleotide binding | 4894 | 208 | 152.8 | 3.00E-07 |
| GO:1901265 | nucleoside phosphate binding | 4895 | 208 | 152.83 | 3.10E-07 |
| GO:0016829 | lyase activity | 900 | 57 | 28.1 | 3.40E-07 |
| GO:0032555 | purine ribonucleotide binding | 4137 | 181 | 129.17 | 3.90E-07 |
| GO:0046527 | glucosyltransferase activity | 364 | 31 | 11.36 | 4.90E-07 |
| GO:0032553 | ribonucleotide binding | 4189 | 182 | 130.79 | 5.80E-07 |
| GO:0017076 | purine nucleotide binding | 4457 | 191 | 139.16 | 7.00E-07 |
| GO:0016597 | amino acid binding | 90 | 14 | 2.81 | 7.40E-07 |

We then examined genetic variants that reached fixation during the domestication of tomato and potato, to identify those with the strongest effects. These are variants for which variation exists in the wild tomato relatives, (e.g. *S. pennellii*), and thus such mutations could be rescued through breeding. To this goal, we repeated the same procedure used for polymorphic variants for fixed mutations evolved in tomato and potato with respect to their wild ancestors (defined as for the proxy-neutral set). For tomato, we observed a strong enrichment for genes involved in NAD(P)H dehydrogenase activity and RNA endonuclease activity (Table 5). For potato, the most notable enrichment is that of violaxanthin de-epoxidase activity genes, with 4 genes out of 9 containing at least a highly deleterious mutation compared to the expected 0.2 (40x fold enrichment) (Table 6). Interestingly, this pathway has been shown to be involved in tuber yield and CO_2_ accumulation^24^. We also repeated the same analysis for *Arabidopsis thaliana*, which differently than tomato and potato did not undergo domestication. For this species, observed an enrichment for genes involved in gene and epigenomic regulation, including methyl-CpNpG binding genes, helicase activity and histone H3 methyltransferase activity (Table 7).

**Table 5.**
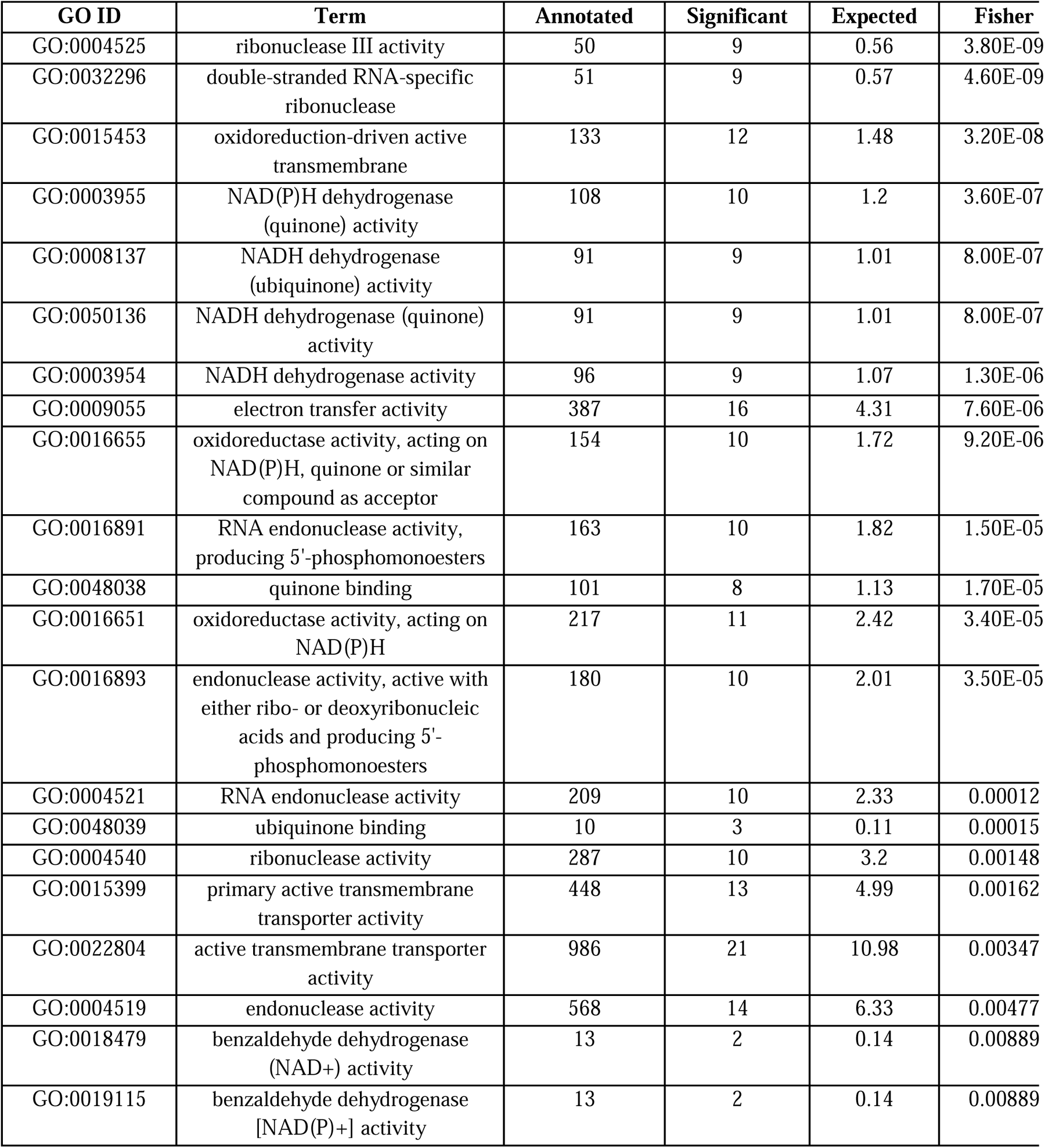
Top 20 gene ontology categories for the genes carrying the most deleterious fixed variants in tomato. The columns describe the GO ID (GO ID), a brief description of the ontology (Term), the total number of annotated (Annotated), significant (Significant) and expected (Expected) hits for each specific ontology, and the p-value (Fisher). Higher Significant than Expected indicates an enrichment.

**Table 6.** Top 20 gene ontology categories for the genes carrying the most deleterious fixed variants in potato. The columns describe the GO ID (GO.ID), a brief description of the ontology (Term), the total number of annotated (Annotated), significant (Significant) and expected (Expected) hits for each specific ontology, and the p-value (Fisher). Higher Significant than Expected indicates an enrichment.

| **GO.ID** | **Term** | **Annotated** | **Significant** | **Expected** | **Fisher** |
| --- | --- | --- | --- | --- | --- |
| GO:0035639 | purine ribonucleoside triphosphate bindi... | 3631 | 122 | 81.17 | 1.80E-06 |
| GO:0005524 | ATP binding | 3215 | 108 | 71.87 | 8.20E-06 |
| GO:0000166 | nucleotide binding | 4697 | 145 | 105 | 1.60E-05 |
| GO:1901265 | nucleoside phosphate binding | 4697 | 145 | 105 | 1.60E-05 |
| GO:0046422 | violaxanthin de-epoxidase activity | 9 | 4 | 0.2 | 2.80E-05 |
| GO:0036094 | small molecule binding | 4957 | 150 | 110.82 | 3.10E-05 |
| GO:0003824 | catalytic activity | 12450 | 325 | 278.33 | 3.80E-05 |
| GO:0043168 | anion binding | 4730 | 142 | 105.74 | 8.10E-05 |
| GO:0032553 | ribonucleotide binding | 4262 | 130 | 95.28 | 8.90E-05 |
| GO:0097367 | carbohydrate derivative binding | 4316 | 131 | 96.49 | 0.0001 |
| GO:0043167 | ion binding | 7737 | 214 | 172.96 | 0.00011 |
| GO:0017076 | purine nucleotide binding | 4415 | 133 | 98.7 | 0.00013 |
| GO:0032555 | purine ribonucleotide binding | 4206 | 127 | 94.03 | 0.00017 |
| GO:0016491 | oxidoreductase activity | 2794 | 90 | 62.46 | 0.00024 |
| GO:0030976 | thiamine pyrophosphate binding | 15 | 4 | 0.34 | 0.00028 |
| GO:0050997 | quaternary ammonium group binding | 15 | 4 | 0.34 | 0.00028 |
| GO:0022804 | active transmembrane transporter activity | 374 | 20 | 8.36 | 0.00032 |
| GO:0016705 | oxidoreductase activity, acting on paire... | 815 | 34 | 18.22 | 0.00041 |
| GO:0030554 | adenyl nucleotide binding | 3988 | 119 | 89.15 | 0.00047 |
| GO:0032559 | adenyl ribonucleotide binding | 3790 | 113 | 84.73 | 0.00069 |
| GO:0016772 | transferase activity, transferring phosp... | 2428 | 78 | 54.28 | 0.0007 |

**Table 7.** Top 20 gene ontology categories for the genes carrying the most deleterious fixed variants in *A. thaliana* (SL5.0). The columns describe the GO ID (GO.ID), a brief description of the ontology (Term), the total number of annotated (Annotated), significant (Significant) and expected (Expected) hits for each specific ontology, and the p-value (Fisher). Higher Significant than Expected indicates an enrichment.

| GO.ID | Term | Annotated | Significant | Expected | Fisher |
| --- | --- | --- | --- | --- | --- |
| GO:0010428 | methyl-CpNpG binding | 8 | 5 | 0.26 | 1.70E-06 |
| GO:0010429 | methyl-CpNpN binding | 8 | 5 | 0.26 | 1.70E-06 |
| GO:0140098 | catalytic activity, acting on RNA | 516 | 37 | 16.52 | 5.10E-06 |
| GO:0140640 | catalytic activity, acting on a nucleic ... | 748 | 44 | 23.95 | 9.10E-05 |
| GO:0003724 | RNA helicase activity | 94 | 11 | 3.01 | 0.00021 |
| GO:0008186 | ATP-dependent activity, acting on RNA | 94 | 11 | 3.01 | 0.00021 |
| GO:0008327 | methyl-CpG binding | 20 | 5 | 0.64 | 0.00035 |
| GO:0140657 | ATP-dependent activity | 815 | 44 | 26.1 | 0.00058 |
| GO:0004386 | helicase activity | 145 | 13 | 4.64 | 0.00081 |
| GO:0003824 | catalytic activity | 8407 | 312 | 269.22 | 0.00096 |
| GO:0140938 | histone H3 methyltransferase activity | 26 | 5 | 0.83 | 0.00125 |
| GO:0061659 | ubiquitin-like protein ligase activity | 284 | 19 | 9.09 | 0.00216 |
| GO:0140101 | catalytic activity, acting on a tRNA | 128 | 11 | 4.1 | 0.00278 |
| GO:0008670 | 2,4-dienoyl-CoA reductase (NADPH) activity | 3 | 2 | 0.1 | 0.00301 |
| GO:0042054 | histone methyltransferase activity | 49 | 6 | 1.57 | 0.00459 |
| GO:0018024 | histone lysine N-methyltransferase activity | 22 | 4 | 0.7 | 0.00482 |
| GO:0008175 | tRNA methyltransferase activity | 12 | 3 | 0.38 | 0.0058 |
| GO:0140096 | catalytic activity, acting on a protein | 2550 | 104 | 81.66 | 0.00617 |
| GO:0061630 | ubiquitin protein ligase activity | 276 | 17 | 8.84 | 0.00806 |
| GO:0016887 | ATP hydrolysis activity | 466 | 25 | 14.92 | 0.00886 |
| GO:0016760 | cellulose synthase (UDP-forming) activity | 27 | 4 | 0.86 | 0.01019 |

### Characterization of genetic variation in tomato

Finally, we focused on tomato leveraging the estimated scores to identify the most deleterious variants identified in GWAS studies^14^ and the most deleterious variants present in genomic regions identified as experiencing positive selection during the domestication of tomato and involved in selective sweeps^25^.

For GWAS we did not identify any single highly deleterious variant (PHRED-scaled score >30) (Table 8). Notice that the GWAS performed by Zhou^14^, focused on identifying genetic variants associated with secondary metabolites in tomato fruits. It is possible that none of these variants strongly affect fitness. This is in stark contrast with what is observed using similar methods to ours in humans, where GWAS are mostly used to identify disease-causing variants^6^.

We examined the genetic variants found in regions identified as selective sweeps in tomatoes by Razifard et al^25^. In this study the authors segmented selective sweeps as events occurring at different times of tomato domestication: two different routes of northward spread of *S. lycopersicum var. cerasiforme* (Table 9 and Table 10) and the redomestication of *S. lycopersicum var. lycopersicum* (Table 11). We find 5, 13, and 14 highly deleterious variants for these three events, respectively, with a G->A variant for the redomestication selective sweep having a score of 47, thus among the most deleterious mutations identified, found at high frequency (∼20% of tomatoes). This gene affects a PRAM-predicted gene identified as 24043 on chromosome 2. We hypothesize that the variants identified here are promising targets for functional studies and to get insights on the deleterious mutations that accumulated and hitchhiked during the domestication of tomatoes.

We visualized the regions affected by these mutations by showing the estimated effect of all potential mutations for the genes carrying the most deleterious mutations in Tables (3-7) (Figure 7-11). These plots also illustrate how our maps of fitness/deleteriousness effect can be used to explore the landscape of effects once a genomic region is mutated. We see how stop-gain variants are usually with a stronger effect than non-synonymous variants, and in turn non-synonymous with a stronger effect than synonymous ones. For instance, the top polymorphic variant for *A. thaliana,* the variant with the highest predicted deleteriousness score is a stop-gain variant. We also see how mutations occurring in introns usually have a much weaker effect than those occurring in coding regions.

Interestingly, we find a few synonymous mutations among those fixed in tomato, potato and A*. thaliana* with scaled PHRED scores larger than 30. This is consistent with increasing evidence that synonymous mutations can have severe impact on fitness. For instance, synonymous mutations in representative genes decrease the fitness of yeast strains^26^. Similarly, synonymous mutations were shown to have both positive and negative fitness effects in pseudomonas^27^, and a functional role in cancer^28^. Such mutations can have effects on translation and management regulations^28^. In fact, algorithms have been developed to infer synonymous mutations with functional effects. Here we putatively achieve a similar goal, inferring synonymous variants with putative effects and with importance in the evolution of tomato, potato and *A. thaliana*.

It is also important to notice that these are fixed mutations (thus generally more neutral and part of our proxy-neutral set), and thus not the most deleterious variants in the genome: only the most deleterious ones among those present nowadays in all tomatoes, all *A. thaliana* and all potatoes, and with existing variation in wild relative. Hence these mutations are certainly not lethal, and for potatoes and tomatoes, likely neither strongly decreasing fitness in a domesticated environment.

This is consistent with the presence of some mutations with moderate effects - for example acting on translation efficiency or transcript stability - as it could be for synonymous mutations.

**Figure 7.**
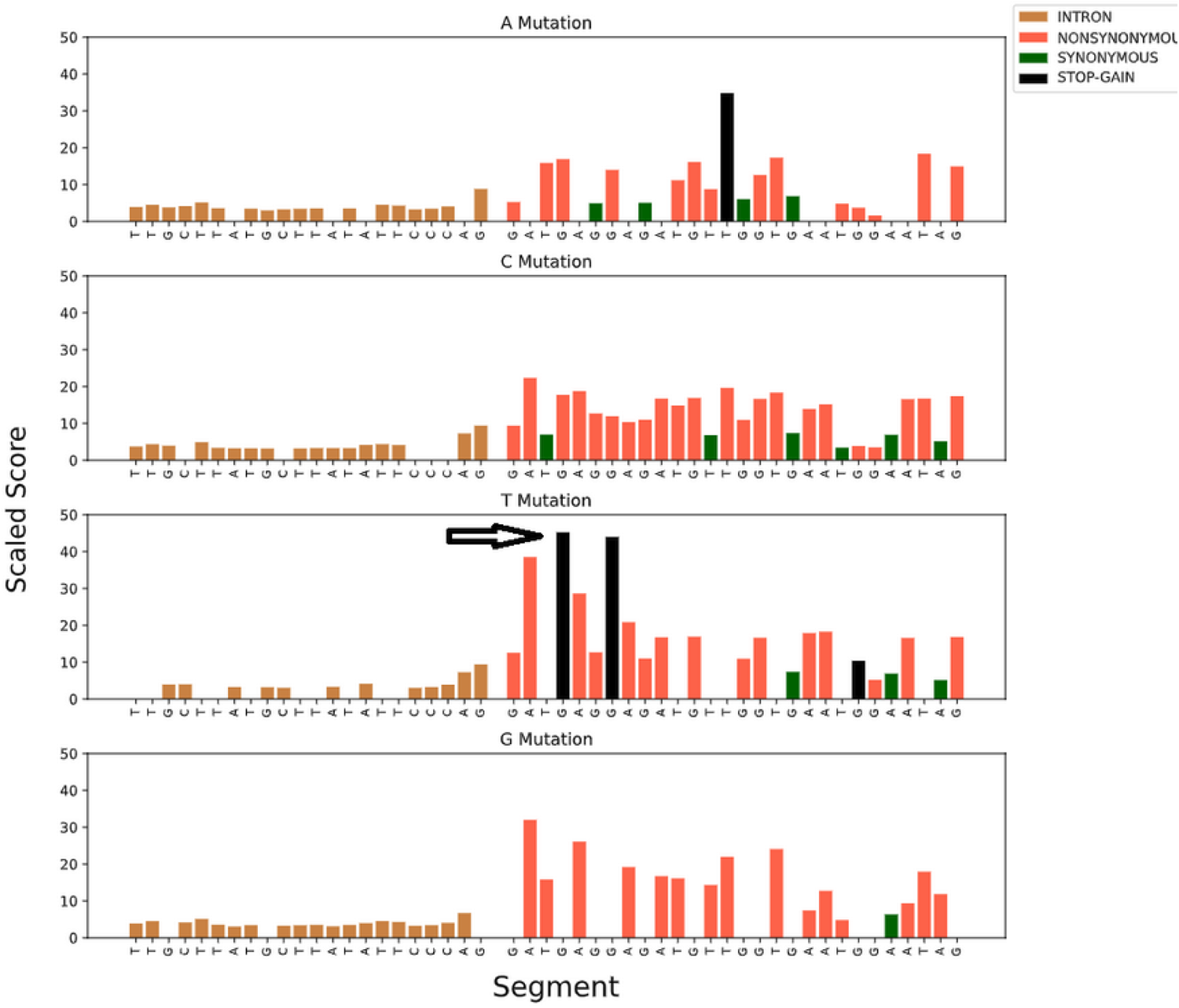
Fitness map spanning 50 bps [2:8695958-8696008] around top scored deleterious variant of polymorphic variants for *A. thaliana*. Each panel presents PHRED-scaled scores for the sequence spanning the variant, considering all positions mutated to a specific allele.

**Figure 8.**
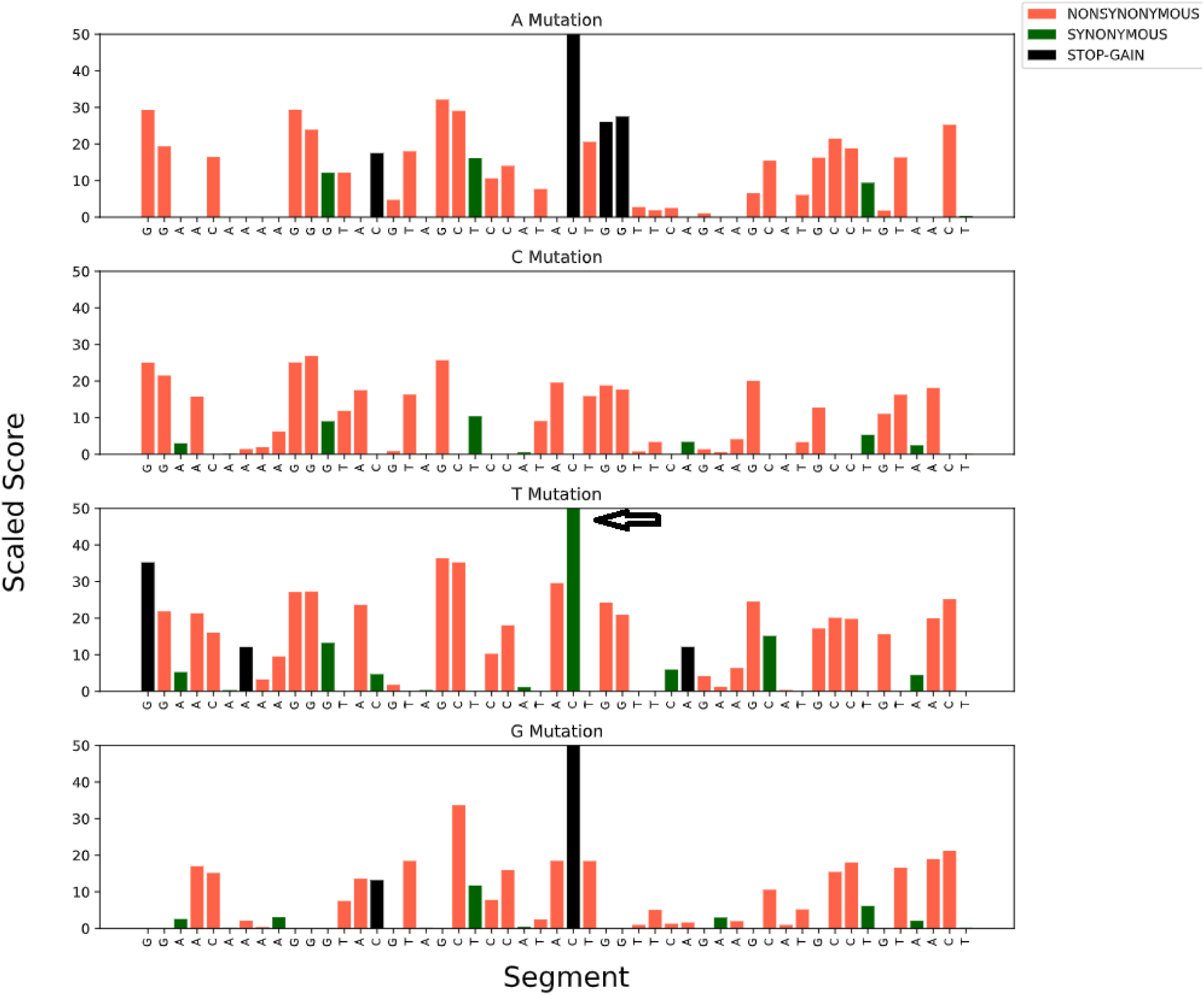
Fitness map spanning 50 bps [1:14606670-14606720] around top scored deleterious variant of polymorphic variants for *S. lycopersicum*. Each panel presents PHRED-scaled scores for the sequence spanning the variant, considering all positions mutated to a specific allele.

**Figure 9.**
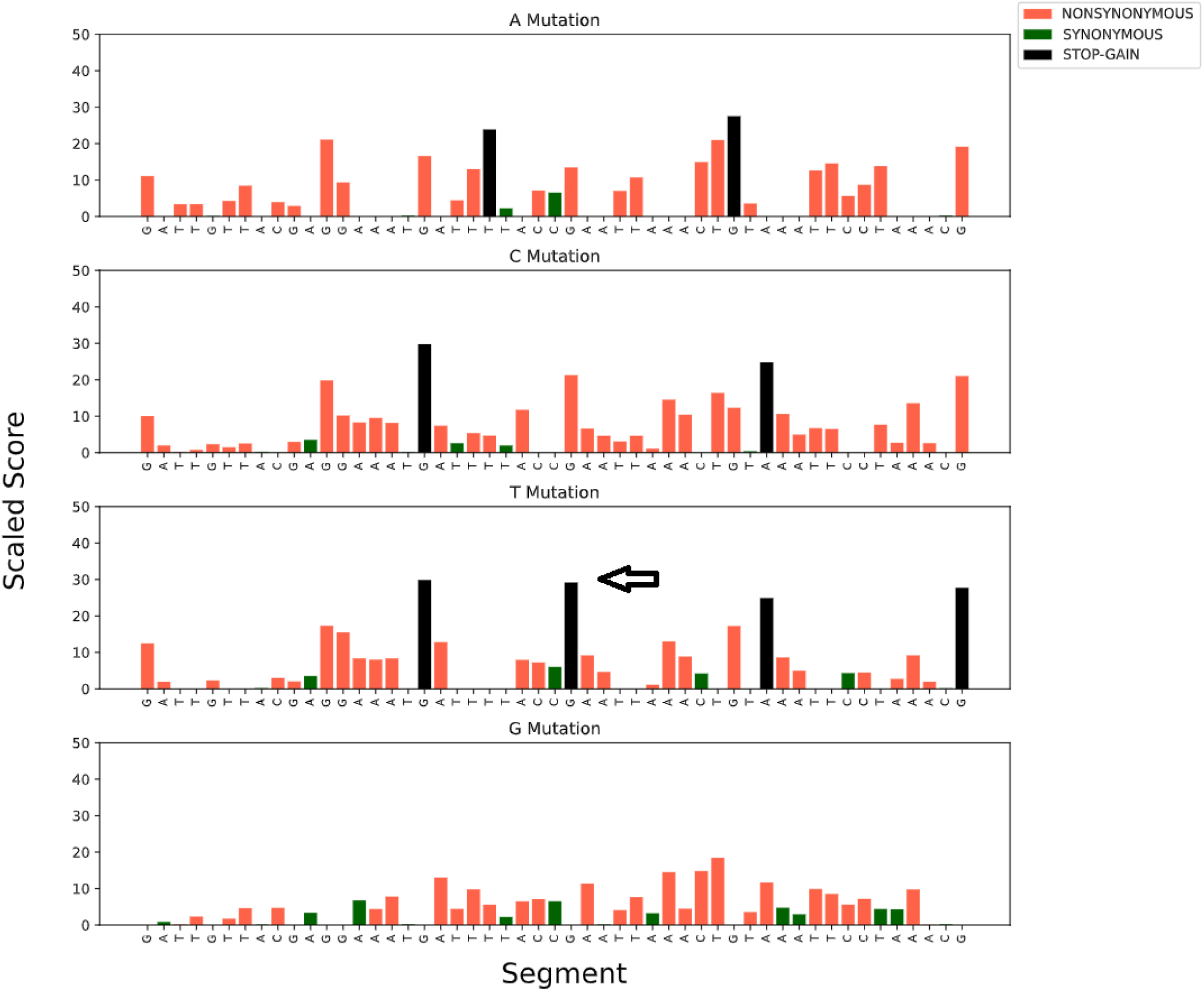
Fitness map spanning 50 bps [4:14392511-14392561] around top scored neutral variant of fixed neutral variants for *A. thaliana*. Each panel presents PHRED-scaled scores for the sequence spanning the variant, considering all positions mutated to a specific allele.

**Figure 10.**
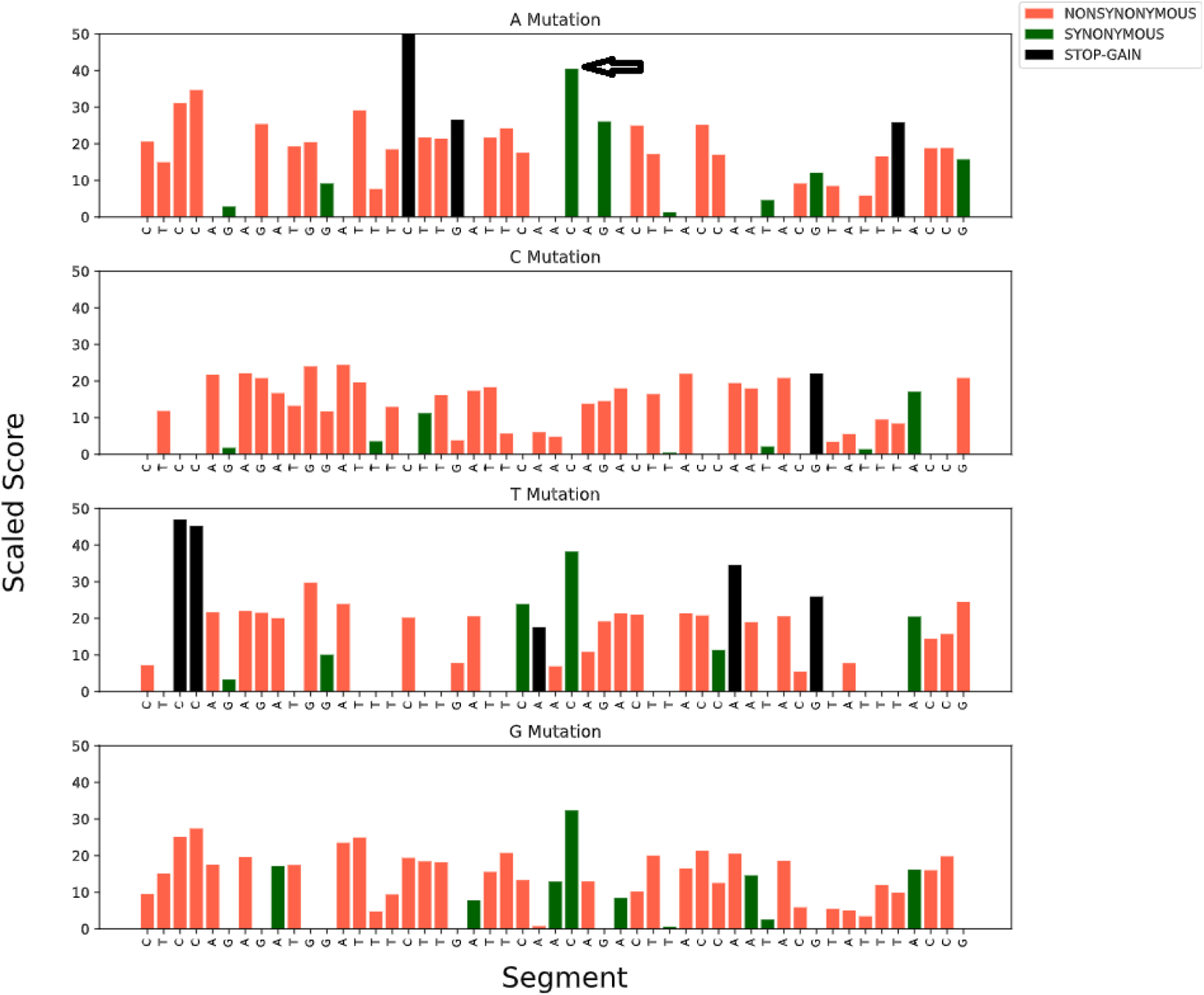
Fitness map spanning 50 bps [12:888705-888755] around top scored neutral variant of fixed neutral variants for *S. lycopersicum*. Each panel presents PHRED-scaled scores for the sequence spanning the variant, considering all positions mutated to a specific allele.

**Figure 11.**
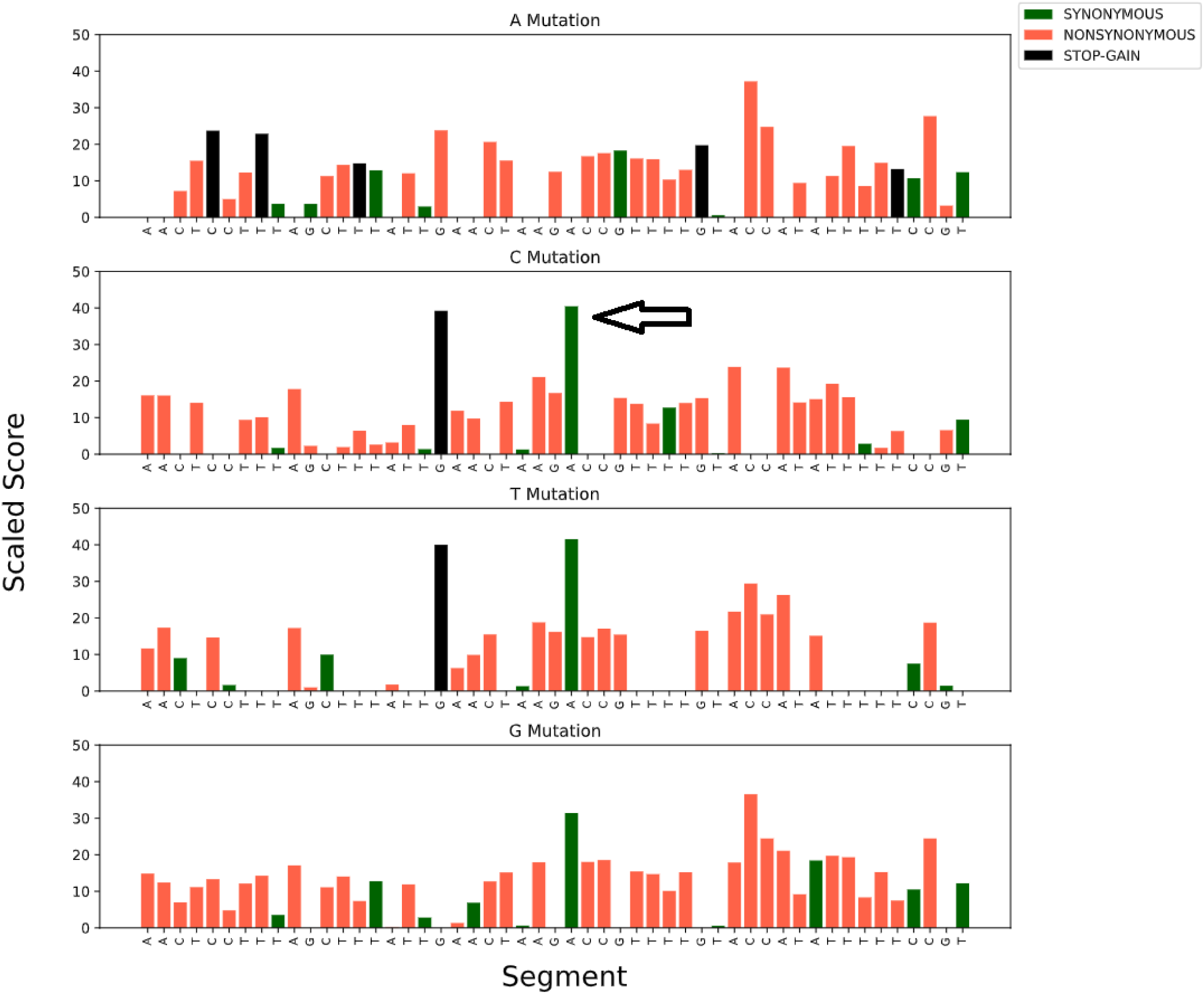
Fitness map spanning 50 bps [5:54298918-54298958] around top scored neutral variant of fixed neutral variants for *S. tuberosum*. Each panel presents PHRED-scaled scores for the sequence spanning the variant, considering all positions mutated to a specific allele.

**Table 8.** Top 30 GWAS variants with the most deleterious effects.

| Chromosome | Start | End | Reference Allele | Alternative Allele | Score | Allele Frequency | Gene ID |
| --- | --- | --- | --- | --- | --- | --- | --- |
| 7 | 3494149 | 3494150 | C | T | 23.3 | 3.584E-01 | Solyc07T000372 |
| 6 | 47441853 | 47441854 | C | A | 21.2 | 2.103E-01 | Solyc06T002144 |
| 1 | 73840981 | 73840982 | T | G | 20.0 | 1.494E-01 | Solyc01T002140 |
| 10 | 65245711 | 65245712 | G | A | 18.8 | 6.232E-02 | Solyc10T002809 |
| 1 | 74024534 | 74024535 | G | A | 18.7 | 1.962E-01 | Solyc01T002158 |
| 6 | 47442678 | 47442679 | G | A | 17.6 | 3.931E-01 | Solyc06T002144 |
| 1 | 74040550 | 74040551 | G | A | 16.9 | 1.643E-01 | - |
| 7 | 3245333 | 3245334 | C | T | 16.3 | 6.317E-01 | Solyc07T000351 |
| 7 | 3307332 | 3307333 | G | A | 16.0 | 5.050E-01 | - |
| 6 | 47442251 | 47442252 | C | A | 15.9 | 2.082E-01 | Solyc06T002144 |
| 7 | 2985122 | 2985123 | C | A | 15.2 | 1.232E-01 | Solyc07T000327 |
| 2 | 34735092 | 34735093 | C | T | 13.6 | 2.967E-01 | - |
| 2 | 34796718 | 34796719 | G | A | 13.0 | 5.460E-01 | - |
| 2 | 34741731 | 34741732 | G | A | 12.6 | 3.024E-01 | - |
| 7 | 3491958 | 3491959 | C | A | 12.6 | 3.116E-01 | - |
| 2 | 34732645 | 34732646 | G | A | 12.5 | 3.541E-01 | - |
| 7 | 3471514 | 3471515 | G | A | 12.4 | 5.496E-01 | - |
| 1 | 73726237 | 73726238 | C | T | 12.1 | 2.698E-01 | Solyc01T002136 |
| 1 | 74059430 | 74059431 | G | T | 12.1 | 1.664E-01 | Solyc01T002160 |
| 7 | 3491634 | 3491635 | G | A | 12.1 | 6.501E-01 | - |
| 1 | 74179138 | 74179139 | C | T | 11.8 | 2.231E-01 | - |
| 7 | 3492119 | 3492120 | C | A | 11.8 | 6.147E-01 | - |
| 2 | 34800075 | 34800076 | C | T | 11.5 | 5.404E-01 | - |
| 7 | 3489877 | 3489878 | G | T | 11.3 | 5.751E-01 | - |
| 7 | 3488939 | 3488940 | G | A | 11.3 | 6.473E-01 | - |
| 7 | 3485297 | 3485298 | G | A | 11.3 | 6.601E-01 | - |
| 7 | 3407350 | 3407351 | G | A | 11.2 | 2.627E-01 | Solyc07T000363 |
| 7 | 3492179 | 3492180 | G | A | 11.0 | 4.016E-01 | - |
| 2 | 34734990 | 34734991 | G | A | 10.7 | 6.246E-01 | - |
| 2 | 34807251 | 34807252 | G | A | 10.7 | 3.173E-01 | - |

**Table 9.**
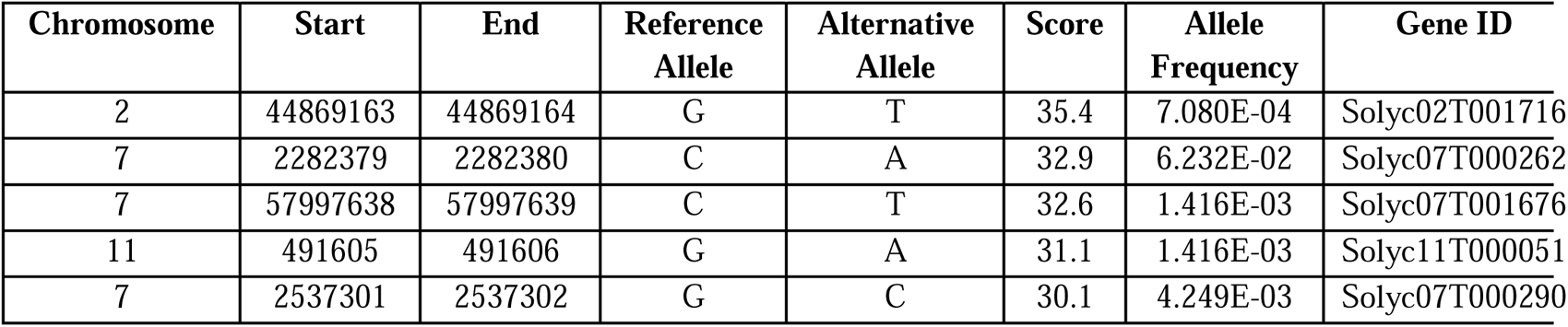
Top deleterious variants with PHRED-scaled score larger than 30 for the first northward spread of *Solanum lycopersicum var cerasiforme* (denoted as North 1 in Razifard et al.).

**Table 10.**
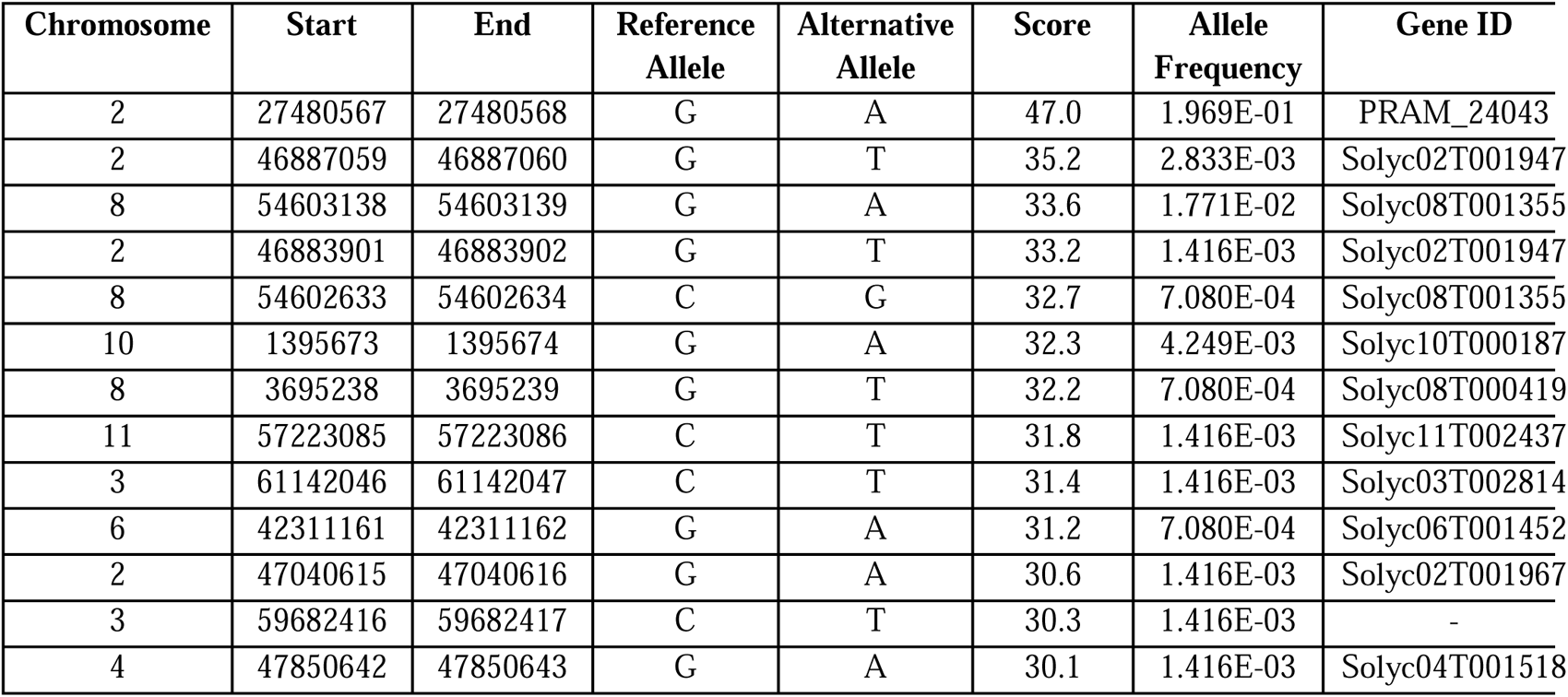
Top deleterious variants with PHRED-scaled score larger than 30 for the second northward spread of *Solanum lycopersicum var cerasiforme* (denoted as North 2 in Razifard et al.).

**Table 11.** Top deleterious variants with PHRED-scaled score larger than 30 for redomestication event of *Solanum lycopersicum var lycopersicum*.

| Chromosome | Start | End | Reference Allele | Alternative Allele | Score | Allele Frequency | Gene ID |
| --- | --- | --- | --- | --- | --- | --- | --- |
| 4 | 55647577 | 55647578 | C | A | 37.7 | 7.080E-04 | Solyc04T001943 |
| 4 | 53594388 | 53594389 | C | A | 37.4 | 7.082E-03 | Solyc04T001815 |
| 4 | 53594572 | 53594573 | C | T | 35.2 | 2.833E-03 | Solyc04T001815 |
| 4 | 54524310 | 54524311 | G | A | 33.3 | 7.082E-03 | Solyc04T001866 |
| 4 | 53594520 | 53594521 | C | G | 33.2 | 7.080E-04 | Solyc04T001815 |
| 9 | 5828620 | 5828621 | C | G | 32.5 | 7.082E-03 | Solyc09T000648 |
| 9 | 5828514 | 5828515 | G | A | 32.1 | 1.416E-03 | Solyc09T000648 |
| 9 | 5828222 | 5828223 | G | A | 31.9 | 1.416E-03 | Solyc09T000648 |
| 4 | 53460216 | 53460217 | C | A | 31.6 | 6.870E-02 | Solyc04T001810 |
| 3 | 63892157 | 63892158 | G | A | 31.4 | 1.416E-03 | Solyc03T003206 |
| 3 | 63895396 | 63895397 | C | T | 30.7 | 4.249E-03 | - |
| 1 | 83904540 | 83904541 | G | A | 30.4 | 4.958E-03 | Solyc01T003223 |
| 9 | 65255249 | 65255250 | G | T | 30.2 | 2.833E-03 | Solyc09T002396 |
| 4 | 55663423 | 55663424 | C | A | 30.1 | 7.080E-04 | - |

## Discussion/Conclusion

In this study we built maps of fitness effects for three different species. We successfully applied, for the first time, a multi-omics approach to study deleterious variation in plants. This approach provides good correlation with allele frequencies, indicating that it can complement and even outperform existing methods to characterize the deleterious load in crops. We showed that the current availability of epigenomic, transcriptomic and population data for many plant species already allows researchers to apply this approach widely. Remarkably, we were able to apply this method successfully both to *A. thaliana*, a model species for which a large abundance of data exists, and to crops for which epigenomic and transcriptomic data are more (tomato, 8 epigenetic marks; population data constituted by 332 individuals fully sequenced) or less (potato, 0 epigenetic marks; non-significant population data) abundant. Noticeably, we trained out models without using population data to establish which mutations were truly fixed. This is a modification that we implemented in respect to the original CADD paper^6^, motivated by heuristic and practical reasons. First, we noticed that for both Arabidopsis and tomato, we could obtain a better proxy-neutral set without requiring the presence of this data. Secondly, we reasoned that for many applications in non-model organisms and in many crop species, comprehensive population data are often missing, scarce, or possibly introducing artifacts. Thus, we managed to build multi-omic fitness maps using annotations, transcriptomic and a limited number of epigenomic marks not only for *A. thaliana*, for which such data abound, but also for two crops with very different quantities and quality of omics data. By doing this, we show that this approach could in principle be applied to many more species, and an exciting perspective is to extend the pipeline designed here to more crops, and even to the entire NCBI, by automation. In fact, many features used for the training of our model can be extracted in an automated way from the genome sequence, genomic annotations and programs that generate conservation scores (in this study leveraging the *msa* pipeline^7^). The more time-consuming step is that of manually collecting epigenomic and other experimental features - a procedure that highly depends on the data available for the species, and thus whose automation is particularly challenging. However, we designed specific rules that can be applied quite generally, and potentially implemented in a semi-automated pipeline, where species-specific repositories are specified by the researcher, while the feature aggregation, creation of the training datasets of mutations and the training are automated.

To show the applicability of this method, we also used the deleteriousness scores computed here to study the effect of genetic variation in the three species, with a focus on tomato and on the fitness effects of mutations which arose during its domestication and genetic variation observed nowadays in tomato populations. We identified the most deleterious mutations currently segregating in tomatoes, and those that reached fixation during evolution. Such mutations putatively represent deleterious mutations that underwent fixation because of drift or genetic hitchhiking, phenomena that are particularly strong during domestication. However, it is important to emphasize that such mutations could represent highly functional ones that were deleterious in the wild - or that share features of mutations that have been deleterious through tomato history - but that are actually advantageous or neutral under farming conditions. In fact, domestication involves strong changes in the environment and in the type of selective pressures that domesticated organisms experience. This highlights both the usefulness and the limitation of methods to infer maps of fitness effects in domesticated species: such methods have the potential to identify genetic deleterious variation which could be used for breeding or gene editing purposes; however, they might fail to unambiguously identify deleterious mutations, and instead they could point to mutations which are highly functional.

## Methods

### Dataset construction

The creation of the dataset was based on a fasta file for each genome. Using Biopython^29^ the fasta file was parsed into a bed file containing 1 bp per row. Onto this file, multiple features were mapped or intersected using bedtools^30^. First, ‘base’ features were generated – these were extract from gene annotations or from the fasta file. Namely, codon position, gene distance, exon and gene counts, and genomic regions were extracted from annotations using python’s pandas^31^ package. Depending on the species, other annotations were also used, such as repeats and splice regions. A measurement for sequence uniqueness, which we name ‘kmers’, was extracted from the fasta file using gem3-mapper^32^ and further parsed to bed format using bedops^33^. The window size around the position was chosen based on the distribution of the final score. We opted to make it as expressive as possible. GC content for a given window was extracted from the fasta using bedtools.

Next, conservation scores were added. These were generated following the *msa* pipeline^7^. This pipeline was used to create pairwise alignment to our reference species [first rule, ‘align’] using default parameters. Next we used multiz’ roast^34^ to create a single multiple alignment format [.maf] file. A tree was generated for this step using mashtree^35^ and the species genomes [fasta format]. As described in the Results seciton, some species had to be removed from this multiple alignment file, using mafTools’^36^ mafFilter. The reference species was masked using awk. The filtered file was then used again for the msa pipeline’s last rule [‘call_conservation’] to generate the following conservation scores: GERP++^4^, PhyloP^5^, PhastCons^5^. For GERP++ and PhyloP, the default parameters of the *msa* pipeline were used. For PhastCons, we allowed the model to estimate the parameters.

We added amino acid substitution information using SIFT^37^, and two confusion matrices: BLOSUM62^38^ and grantham^39^. These were added to our training set using python in a parsing step further down the line. For SIFT, we built a database per species using SIFT’s perl scripts to do so.

Lastly, experimental features were added. Whenever raw data was used, the papers’ methods were replicated, if specified. This section varies per species:

For A. thaliana, we used PCSD^40^. Besides using their final chromatin states mapping of the genome, we also used their epigenetic markers data, including methylation, histone modifications, chromatin associated factors, DNA accessibility, and transcription factors. For the last three, we aggregated all of their samples per modification by maximum value, and substituted zero for missing positions in the genome. For expression, we used normalized counts deposited into NCBI41 [GSE80744].

For *S. lycopersicum*:

- Expression data was taken from solomics^22^, median and precision were extracted using python.
- For histone modifications, we used:

`o` H3K27ME3, H3K4ME3, H3K9Ac^42^. Each timestamp (0da, 4dpa, 4iaa) was parsed individually, and then combined aggregating for a maximum value. Data was aligned to SL5.0 using bowtie2^43^, with both replicates. Samtools^44^ was used to clean duplicates via rmdup. Peaks were called using MACS2^45^ with parameters -g 8.15e8 –pvalue 1e-5.
`o` H3K27ac^46^ was parsed as described by the paper.
- Methylome using BS-seq^46^ was parsed as described by the paper.
- DNAse1 hypersensitivity sites^47^ – both samples were parsed individually and then combined aggregating for a maximum value. Per sample, we used bowtie2 to align to SL5.0, with only 1 replicate. Next, duplicates were cleaned using samtools rmdup, and peaks were called using MACS2 with parameters --nomodel --broad --broad-cutoff 0.01 -g 8.15e8.
- Regulatory data^48^ was lifted over to SL5.0 coordinates from SL2.4 using HAL^49^ package’s halLiftover. The alignment file used was generated using the *msa* pipeline^7^, and then converted using HAL’s maf2hal. Further parsing was done using bedtools and pandas.
- Chromatin accessibility data^50^ was parsed following the paper’s procedure.
- Short RNAs^51–53^ were obtained via a few papers. We obtained miRNA, circRNA and lncRNA positions, and all were lifted from SL3.0 following the lifting procedure described above. For miRNA we had a few sources and thus aggregated using the maximum value.

From all these epigenetic markers, we used ChromHMM54 to build a map of chromatin states along the genomes. We used a bin size of 40 and a prediction of 8 states. This was done solely for *S. lycopersicum*, as we already had the ChromHMM map for Arabidopsis [via PCSD], and we did not use any epigenetic markers for *S. tuberosum*.

The dataset was thus complete. For positions coding for genes, we had 4 possible alleles – they were replicated fourfold.

### Dataset Labeling

We divided our training dataset into two labels – proxy-neutral and proxy-deleterious. The proxy neutral was ascertained as described in the main text, excluding gaps. First, the *msa* pipeline^7^ output file in MAF format was parsed in awk to obtain an intermediate BED-format table containing all positions in the genome. This was used to filter out genomic positions without at least 13 aligned species and all further operations. The proxy-deleterious was generated by selecting positions at which the reference genome differs from the species used as outgroup (*S. pennellii* for tomato, *A. arenosa* for *A. thaliana* and *S. okadae* for potato) and randomly sampling a new position following a discretized Gaussian with standard deviation 10kbp and excluding multiple mutations occurring in the same position as previously sampled ones, or mutations occurring in regions of the genome not analyzed as excluded by the filters (i.e. less than 13 species aligned). Each new mutation retained the type of substitution, e.g. from A to C, in order to keep the mutation spectrum as close as possible to that of newly arising mutations. Since the positions of new mutations are permuted around the originally observed ones, this scheme aims to maintain the same local mutation rates, while generating new positions for mutations which are unconstrained to signal of selection – which we expect to vary on an exon-like scale or shorter, i.e. 200bp range. Note that this scheme is different from that implemented in previous implementation of Kircher et al. 2014 and previous iterations of this approach, that instead relied on mutation rates estimated via phylogenetic models. We opted for a simpler permutation-based approach, which is independent from local rates estimated via phylogenetic models, since estimating local phylogenetic relationships and mutation rates based on often patchy multiple whole genome alignments is often challenging for plants, in particular domesticated ones, and this could lead to unforeseen biases in overall or local mutation rates. These operations were performed using custom awk and R scripts.

Following generating these positions, we crossed them with our dataset. Positions in which the number of species aligned to was lower than 13 were removed. A few more features were added, including whether the mutation is a transition/transversion, and the confusion matrices scores. Categorical features were 1-hot encoded. Finally, we normalized our features independently, using standardization (z-score scaling) using scikit learn’s scaler^55^. We did not include indels or hard masked regions in our training dataset. We used a 1:1 ratio for our labels, while proxy-neutral mutations were always the limiting factor.

This set went through further parsing, adding protein features (such as substitution matrices scores), one hot encoding for categorical features and finally normalization (z-scaling).

### Model Training

After comparing a naïve NN model to that of Kircher’s^6^, we trained an ensemble of 50 SVM learners. The training set was bootstrapped 50 times and fed into 50 SVM learners with a linear kernel. The learning was done in batches of size 8,192. The predictions were aggregated using mean. The prediction scores were in the range [0,1], 0 noting neutral variants, and 1 noting deleterious ones. In order to scale our predictions scores to a PHRED-like scale, we generated quantiles using our entire proxy-deleterious set per species.

The naïve neural network, implemented with pytorch^56^. The network was a simple feedforward comprised of two linear layers, with ReLU activation in between and a sigmoid at the output.

Following are the features used for the models described:

#### Solanum lycopersicum

**Table 12.** Features used for the SVM model for *S. lycopersicum*.

| Feature | Description | Type | Values |
| --- | --- | --- | --- |
| ref_nucleotide | reference nucleotide for position | discrete | A/C/T/G |
| kmers_20 | value for uniqueness of sequence of size 20 around position | continuous | - |
| gc_content_35_window | gc content in a 35 window around position | continuous | - |
| gene_overlap_count | overlapping genes count for position | continuous | - |
| exon_overlap_count | overlapping exons count for position | continuous | - |
| repeats_class | whether position is of a repeat class | discrete | DNA/LTR/OTHER |
| repeats_family | whether position is of a repeat family | discrete | Copia/DTC/Gypsy/Helitron/ltrunkown/OTHER |
| codon_pos | codon position for position | discrete | 1/2/3/-1 |
| genomic_region | the genomic region the position is a part of | discrete | CDS/exon/5UTR/gene/mRNA/3UTR/intergenic |
| exp_median | expression normalized counts median for position | continuous | - |
| exp_pre | expression normalized counts precision for position | continuous | - |
| bs_seq | methylation ratio for position | continuous | - |
| atac_seq | peaks for atac-seq | continuous | - |
| dnase1 | peaks for dnase1 hypersensitivity | continuous | - |
| H3K27ac | peaks for h3k27ac chipseq | continuous | - |
| H3K9Ac | peaks for h3k9ac chipseq | continuous | - |
| H3K4ME3 | peaks for h3k4me3 chipseq | continuous | - |
| H3K27ME3 | peaks for h3k27me3 chipseq | continuous | - |
| mi_RNA | whether position is a mi_RNA | discrete | 0/1 |
| lnc_RNA | whether position is a lnc_RNA | discrete | 0/1 |
| circ_RNA | whether position is a circ_RNA | discrete | 0/1 |
| funTFBS | whether position is a functional transcription factor binding site | discrete | 0/1 |
| CEmotif | whether position is a conserved element | discrete | 0/1 |
| motifTFBS | whether position is a transcription factor binding site motif | discrete | 0/1 |
| chromhmm | chromatin state for bin size 40 and 8 states predicted using chromhmm | discrete | 1-8 |
| alt_allele | mutation allele for coding regions | discrete | A/C/T/G |
| variant_type | type of mutation | discrete | synonymous/nonsynonymous/start loss/stop loss/stop gain |
| amino_pos | position of amino acid affected by mutation | continuous | - |
| sift_score | score for mutation | continuous | - |
| sift_median | median score for mutation | continuous | - |
| num_seqs | # sequences in position | continuous | - |
| sift_prediction | final SIFT prediction for mutation | discrete | tolerated/deleterious |
| upstream_dist | upstream distance from closest gene (start or end) | continuous | - |
| downstream_dist | downstream distance from closest gene (start or end) | continuous | - |
| phylop_lrt | phylop score (method LRT) | continuous | - |
| phylop_gerp | phylop score (method GERP) | continuous | - |
| phylop_score | phylop score (method SCORE) | continuous | - |
| gerp_exp | gerp expected substitutions score | continuous | - |
| gerp_rej | gerp rejected substitutions score | continuous | - |
| gerp_ta | taxa aligned count for alignment | continuous | - |
| phast_est | phastcons score with parameters estimated | continuous | - |

#### Arabidopsis thaliana

**Table 13.** Features used for the SVM model for *A. thaliana*.

| Feature | Description | Type | Values |
| --- | --- | --- | --- |
| ref_nucleotide | reference nucleotide for position | discrete | A/C/T/G |
| kmers_13 | value for uniqueness of sequence of size 13<br>around position | continuous | - |
| gc_content_35_window | gc content in a 35 window around position | continuous | - |
| gene_overlap_count | overlapping gene count for position | continuous | - |
| exon_overlap_count | overlapping exons count for position | continuous | - |
| splice_junction | whether position if part of a splice junction | discrete | 0/1 |
| repeats_class | whether position is a repeat | discrete | 0/1 |
| codon_pos | codon position for position | discrete | 1/2/3/-1 |
| genomic_region | the genomic region the position is a part of | discrete | CDS/exon/5UTR/gene/m<br>RNA/mRNA_TE_gene/p<br>seudogene/transposable_<br>element_gene/shortRNA/<br>3UTR/intergenic |
| exp_median | expression normalized counts median for position | continuous | - |
| methylation_peaks | methylation peaks for methylation markers | continuous | - |
| atacseq_peaks | peaks for atac-seq | continuous | - |
| dnase1 | peaks for dnase1 hypersensitivity markers | continuous | - |
| H3K27Ac | peaks for h3k27ac chipseq | continuous | - |
| H3K9Ac | peaks for h3k9ac chipseq | continuous | - |
| H3K4ME3 | peaks for h3k4me3 chipseq | continuous | - |
| H3K27ME3 | peaks for h3k27me3 chipseq | continuous | - |
| H3K36Ac | peaks for h3k36ac chipseq | continuous | - |
| H3K9ME1 | peaks for h3k9me1 chipseq | continuous | - |
| H3K23Ac | peaks for h3k23Ac chipseq | continuous | - |
| H3K4ME1 | peaks for h3k4me1 chipseq | continuous | - |
| H3K4ME2 | peaks for h3k4me2 chipseq | continuous | - |
| TF_peaks | peaks for transcription factor markers | continuous | - |
| chrom_acc_peaks | peaks for chromatin accessibility markers | continuous | - |
| chromhmm | chromatin state for bin size 200 and 36 states predicted using chromhmm | discrete | 1-36 |
| alt_allele | mutation allele for coding regions | discrete | A/C/T/G |
| variant_type | type of mutation | discrete | synonymous/nonsynonymous/start loss/stop loss/stop gain |
| amino_pos | position of amino acid affected by mutation | continuous | - |
| sift_score | score for mutation | continuous | - |
| sift_median | median score for mutation | continuous | - |
| num_seqs | # sequences in position | continuous | - |
| sift_prediction | final SIFT prediction for mutation | discrete | tolerated/deleterious |
| upstream_dist | upstream distance from closest gene (start or end) | continuous | - |
| downstream_dist | downstream distance from closest gene (start or end) | continuous | - |
| phylop_lrt | phylop score (method LRT) | continuous | - |
| phylop_gerp | phylop score (method GERP) | continuous | - |
| phylop_score | phylop score (method SCORE) | continuous | - |
| gerp_exp | gerp expected substitutions score | continuous | - |
| gerp_rej | gerp rejected substitutions score | continuous | - |
| gerp_ta | taxa aligned count for alignment | continuous | - |
| phast_est | phastcons score with parameters estimated | continuous | - |

#### Solanum tuberosum

**Table 14.**
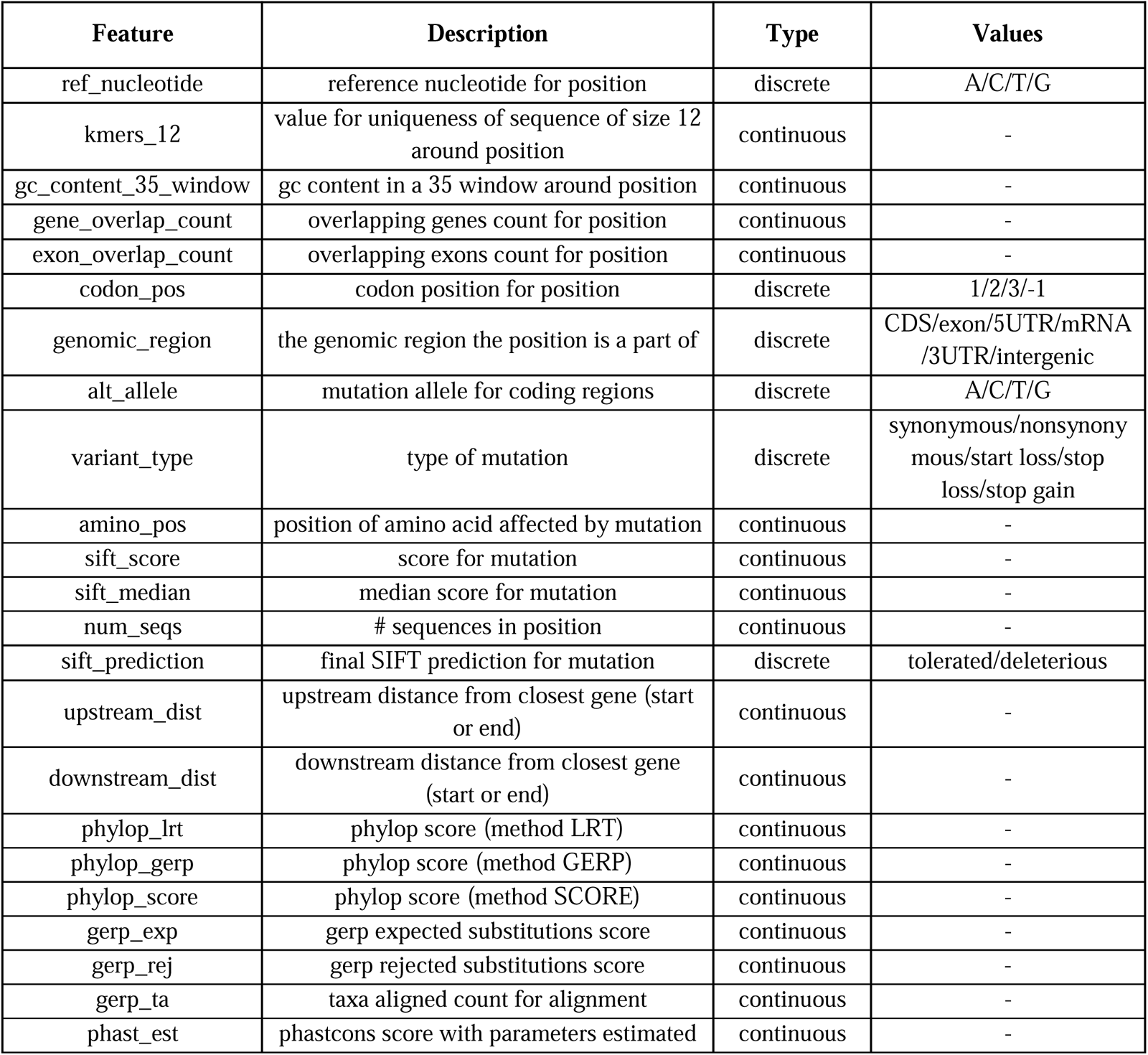
Features used for the SVM model for *S. tuberosum*.

## Code Availability

The code is available at https://github.com/fabrimafe/plantCADD.

## Appendix

**Supplementary Table 1.** Genomes for *S. lycopersicum* alignment.

| Species | Link |
| --- | --- |
| *Anisodus acutangulus* | https://www.ncbi.nlm.nih.gov/datasets/genome/GCA_029783415.1/ |
| *Arabidopsis arenosa* | https://www.ncbi.nlm.nih.gov/datasets/genome/GCA_905216605.1/ |
| *Arabidopsis suecica* | https://www.ncbi.nlm.nih.gov/data-hub/genome/GCA_019202805.1/ |
| *Arabidopsis thaliana 1* | https://www.ncbi.nlm.nih.gov/data-hub/genome/GCF_000001735.4/ |
| *Arabis alpina* | https://www.ncbi.nlm.nih.gov/data-hub/genome/GCA_900128785.1/ |
| *Arabis montbretiana* | https://www.ncbi.nlm.nih.gov/data-hub/genome/GCA_001484125.2/ |
| *Brassica carinata* | https://www.ncbi.nlm.nih.gov/data-hub/genome/GCA_016771965.1/ |
| *Brassica juncea* | https://www.ncbi.nlm.nih.gov/data-hub/genome/GCA_018703725.1/ |
| *Brassica napus* | https://www.ncbi.nlm.nih.gov/data-hub/genome/GCF_020379485.1/ |
| *Brassica nigra* | https://www.ncbi.nlm.nih.gov/data-hub/genome/GCA_016432835.1/ |
| *Brassica oleracea var. oleracea* | https://www.ncbi.nlm.nih.gov/data-hub/genome/GCF_000695525.1/ |
| *Brassica rapa* | https://www.ncbi.nlm.nih.gov/data-hub/genome/GCF_000309985.2/ |
| *Camelina hispida* | https://www.ncbi.nlm.nih.gov/data-hub/genome/GCA_023657505.1/ |
| *Camelina laxa* | https://www.ncbi.nlm.nih.gov/data-hub/genome/GCA_024034495.1/ |
| *Camelina neglecta* | https://www.ncbi.nlm.nih.gov/data-hub/genome/GCA_023864065.1/ |
| *Camelina sativa* | https://www.ncbi.nlm.nih.gov/data-hub/genome/GCF_000633955.1/ |
| *Capsicum annum* | https://www.ncbi.nlm.nih.gov/data-hub/genome/GCF_002878395.1/ |
| *Capsicum baccatum* | https://www.ncbi.nlm.nih.gov/data-hub/genome/GCA_002271885.2/ |
| *Capsicum chinense* | https://www.ncbi.nlm.nih.gov/data-hub/genome/GCA_002271895.2/ |
| *Cucumis melo* | https://www.ncbi.nlm.nih.gov/datasets/genome/GCF_025177605.1/ |
| *Daucus carota* | https://www.ncbi.nlm.nih.gov/datasets/genome/GCA_030127425.1/ |
| *Erysimum cheiranthoides* | https://www.ncbi.nlm.nih.gov/data-hub/genome/GCA_011420285.1/ |
| *Euphorbia peplus* | https://www.ncbi.nlm.nih.gov/datasets/genome/GCA_028411795.1/ |
| *Gossypium herbaceum* | https://www.ncbi.nlm.nih.gov/datasets/genome/GCA_025502765.1/ |
| *Isatis tinctoria* | https://www.ncbi.nlm.nih.gov/data-hub/genome/GCA_010577795.1/ |
| *Lycium barbarum* | https://www.ncbi.nlm.nih.gov/data-hub/genome/GCA_019175385.1/ |
| *Lycium ferocissimum* | https://www.ncbi.nlm.nih.gov/datasets/genome/GCA_029784015.1/ |
| *Nicotiana attenuata* | https://www.ncbi.nlm.nih.gov/data-hub/genome/GCF_001879085.1/ |
| *Oryza sativa Indica Group* | https://www.ncbi.nlm.nih.gov/datasets/genome/GCA_001623345.3/ |
| *Przewalskia tangutica* | https://www.ncbi.nlm.nih.gov/datasets/genome/GCA_029844215.1/ |
| *Pugionium cornutum* | https://www.ncbi.nlm.nih.gov/data-hub/genome/GCA_018901935.1/ |
| *Rhododendron vialii* | https://www.ncbi.nlm.nih.gov/datasets/genome/GCA_030253575.1/ |
| *Solanum appendiculatum* | https://www.ncbi.nlm.nih.gov/data-hub/genome/GCA_018342035.1/ |
| *Solanum arcanum* | https://www.ncbi.nlm.nih.gov/datasets/genome/GCA_027789145.1/ |
| *Solanum chilense* | https://www.ncbi.nlm.nih.gov/data-hub/genome/GCA_006013705.1/ |
| *Solanum chmielewskii* | https://www.ncbi.nlm.nih.gov/datasets/genome/GCA_027704785.1/ |
| *Solanum clarkiae* | https://www.ncbi.nlm.nih.gov/data-hub/genome/GCA_011800125.1/ |
| *Solanum commersonii* | https://www.ncbi.nlm.nih.gov/data-hub/genome/GCA_018258275.1/ |
| *Solanum corneliomulleri* | https://www.ncbi.nlm.nih.gov/datasets/genome/GCA_027704805.1/ |
| *Solanum dulcamara* | https://www.ncbi.nlm.nih.gov/data-hub/genome/GCA_947179165.1/ |
| *Solanum galapagense* | https://www.ncbi.nlm.nih.gov/data-hub/genome/GCA_025758845.1/ |
| *Solanum habrochaites* | https://www.ncbi.nlm.nih.gov/datasets/genome/GCA_027704765.1/ |
| *Solanum lycopersicoides* | https://www.ncbi.nlm.nih.gov/data-hub/genome/GCA_022817965.1/ |
| *Solanum lycopersicum* 5.0 | http://solomics.agis.org.cn/tomato/ftp/SL5.0_masked_genome/ |
| *Solanum melongena* | https://www.ncbi.nlm.nih.gov/data-hub/genome/GCA_000787875.1/ |
| *Solanum neorickii* | https://www.ncbi.nlm.nih.gov/datasets/genome/GCA_027704865.1/ |
| *Solanum okadae* | https://www.ncbi.nlm.nih.gov/data-hub/genome/GCA_023159065.1/ |
| *Solanum ossicruentum* | https://www.ncbi.nlm.nih.gov/data-hub/genome/GCA_025774065.1/ |
| *Solanum pennellii* | https://www.ncbi.nlm.nih.gov/data-hub/genome/GCF_001406875.1/ |
| *Solanum peruvianum* | https://www.ncbi.nlm.nih.gov/datasets/genome/GCA_027705065.1/ |
| *Solanum pinnatisectum* | https://www.ncbi.nlm.nih.gov/data-hub/genome/GCA_009887355.1/ |
| *Solanum rostratum* | https://www.ncbi.nlm.nih.gov/datasets/genome/GCA_029948455.1/ |
| *Solanum sejunctum* | https://www.ncbi.nlm.nih.gov/data-hub/genome/GCA_025642045.1/ |
| *Solanum sitiens* | https://www.ncbi.nlm.nih.gov/data-hub/genome/GCA_016801875.1/ |
| *Solanum stenotomum* | https://www.ncbi.nlm.nih.gov/data-hub/genome/GCF_019186545.1/ |
| *Solanum tuberosum* (Solyntus) | https://www.ncbi.nlm.nih.gov/data-hub/genome/GCA_015076265.1/ |
| *Solanum verrucosum* | https://www.ncbi.nlm.nih.gov/data-hub/genome/GCF_900185275.1/ |
| *Thlaspi arvense* | https://www.ncbi.nlm.nih.gov/data-hub/genome/GCA_911865555.2/ |

**Supplementary Table 2.**
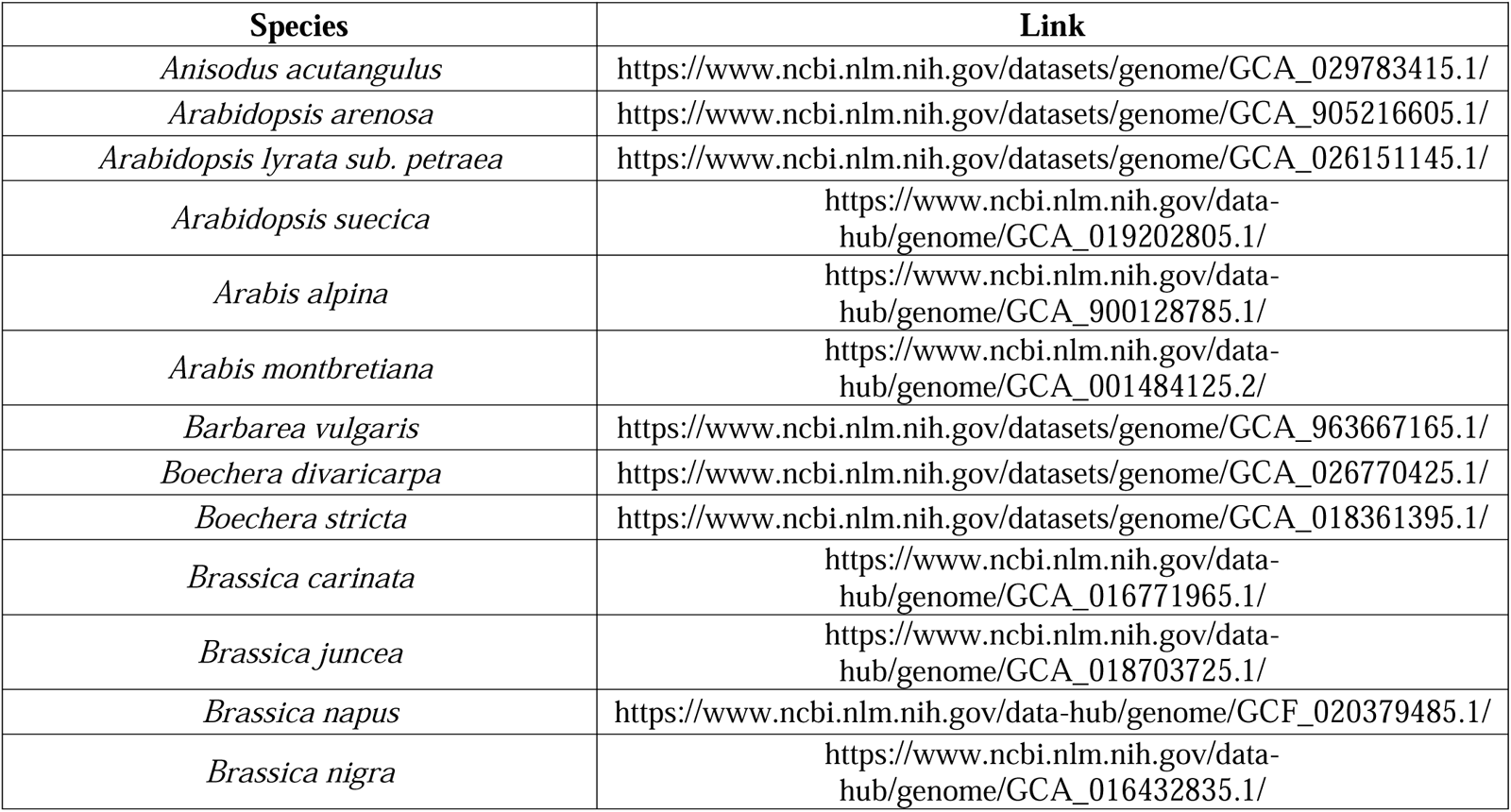
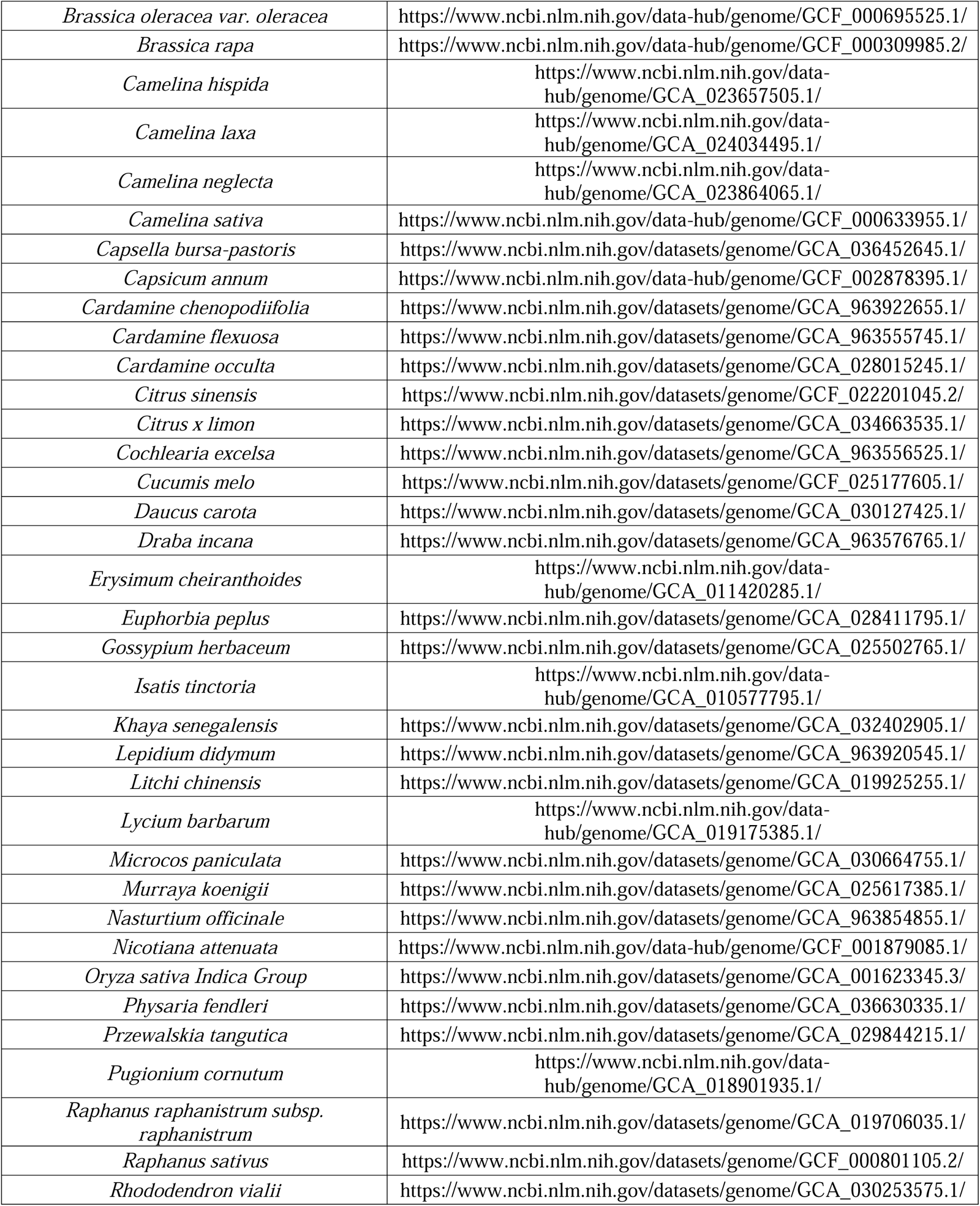
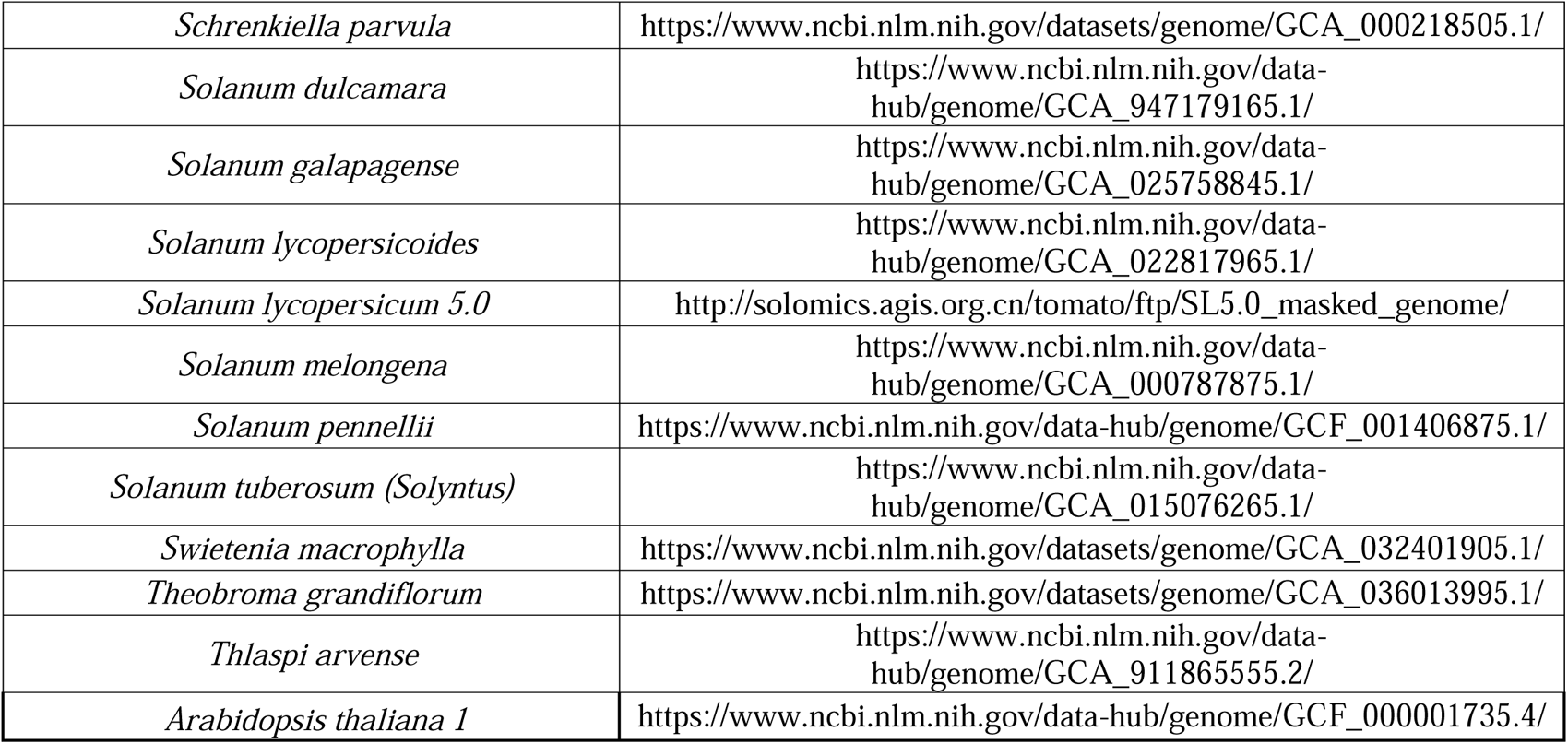
Genomes for *A. thaliana* alignment.

| Species | Link |
| --- | --- |
| *Anisodus acutangulus* | https://www.ncbi.nlm.nih.gov/datasets/genome/GCA_029783415.1/ |
| *Arabidopsis arenosa* | https://www.ncbi.nlm.nih.gov/datasets/genome/GCA_905216605.1/ |
| *Arabidopsis lyrata* sub. *petraea* | https://www.ncbi.nlm.nih.gov/datasets/genome/GCA_026151145.1/ |
| *Arabidopsis suecica* | https://www.ncbi.nlm.nih.gov/data-hub/genome/GCA_019202805.1/ |
| *Arabis alpina* | https://www.ncbi.nlm.nih.gov/data-hub/genome/GCA_900128785.1/ |
| *Arabis montbretiana* | https://www.ncbi.nlm.nih.gov/data-hub/genome/GCA_001484125.2/ |
| *Barbarea vulgaris* | https://www.ncbi.nlm.nih.gov/datasets/genome/GCA_963667165.1/ |
| *Boechera divaricarpa* | https://www.ncbi.nlm.nih.gov/datasets/genome/GCA_026770425.1/ |
| *Boechera stricta* | https://www.ncbi.nlm.nih.gov/datasets/genome/GCA_018361395.1/ |
| *Brassica carinata* | https://www.ncbi.nlm.nih.gov/data-hub/genome/GCA_016771965.1/ |
| *Brassica juncea* | https://www.ncbi.nlm.nih.gov/data-hub/genome/GCA_018703725.1/ |
| *Brassica napus* | https://www.ncbi.nlm.nih.gov/data-hub/genome/GCF_020379485.1/ |
| *Brassica nigra* | https://www.ncbi.nlm.nih.gov/data-hub/genome/GCA_016432835.1/ |
| *Brassica oleracea* var. *oleracea* | https://www.ncbi.nlm.nih.gov/data-hub/genome/GCF_000695525.1/ |
| *Brassica rapa* | https://www.ncbi.nlm.nih.gov/data-hub/genome/GCF_000309985.2/ |
| *Camelina hispida* | https://www.ncbi.nlm.nih.gov/data-hub/genome/GCA_023657505.1/ |
| *Camelina laxa* | https://www.ncbi.nlm.nih.gov/data-hub/genome/GCA_024034495.1/ |
| *Camelina neglecta* | https://www.ncbi.nlm.nih.gov/data-hub/genome/GCA_023864065.1/ |
| *Camelina sativa* | https://www.ncbi.nlm.nih.gov/data-hub/genome/GCF_000633955.1/ |
| *Capsella bursa-pastoris* | https://www.ncbi.nlm.nih.gov/datasets/genome/GCA_036452645.1/ |
| *Capsicum annum* | https://www.ncbi.nlm.nih.gov/data-hub/genome/GCF_002878395.1/ |
| *Cardamine chenopodiifolia* | https://www.ncbi.nlm.nih.gov/datasets/genome/GCA_963922655.1/ |
| *Cardamine flexuosa* | https://www.ncbi.nlm.nih.gov/datasets/genome/GCA_963555745.1/ |
| *Cardamine occulta* | https://www.ncbi.nlm.nih.gov/datasets/genome/GCA_028015245.1/ |
| *Citrus sinensis* | https://www.ncbi.nlm.nih.gov/datasets/genome/GCF_022201045.2/ |
| *Citrus x limon* | https://www.ncbi.nlm.nih.gov/datasets/genome/GCA_034663535.1/ |
| *Cochlearia excelsa* | https://www.ncbi.nlm.nih.gov/datasets/genome/GCA_963556525.1/ |
| *Cucumis melo* | https://www.ncbi.nlm.nih.gov/datasets/genome/GCF_025177605.1/ |
| *Daucus carota* | https://www.ncbi.nlm.nih.gov/datasets/genome/GCA_030127425.1/ |
| *Draba incana* | https://www.ncbi.nlm.nih.gov/datasets/genome/GCA_963576765.1/ |
| *Erysimum cheiranthoides* | https://www.ncbi.nlm.nih.gov/data-hub/genome/GCA_011420285.1/ |
| *Euphorbia peplus* | https://www.ncbi.nlm.nih.gov/datasets/genome/GCA_028411795.1/ |
| *Gossypium herbaceum* | https://www.ncbi.nlm.nih.gov/datasets/genome/GCA_025502765.1/ |
| *Isatis tinctoria* | https://www.ncbi.nlm.nih.gov/data-hub/genome/GCA_010577795.1/ |
| *Khaya senegalensis* | https://www.ncbi.nlm.nih.gov/datasets/genome/GCA_032402905.1/ |
| *Lepidium didymum* | https://www.ncbi.nlm.nih.gov/datasets/genome/GCA_963920545.1/ |
| *Litchi chinensis* | https://www.ncbi.nlm.nih.gov/datasets/genome/GCA_019925255.1/ |
| *Lycium barbarum* | https://www.ncbi.nlm.nih.gov/data-hub/genome/GCA_019175385.1/ |
| *Microcos paniculata* | https://www.ncbi.nlm.nih.gov/datasets/genome/GCA_030664755.1/ |
| *Murraya koenigii* | https://www.ncbi.nlm.nih.gov/datasets/genome/GCA_025617385.1/ |
| *Nasturtium officinale* | https://www.ncbi.nlm.nih.gov/datasets/genome/GCA_963854855.1/ |
| *Nicotiana attenuata* | https://www.ncbi.nlm.nih.gov/data-hub/genome/GCF_001879085.1/ |
| *Oryza sativa Indica Group* | https://www.ncbi.nlm.nih.gov/datasets/genome/GCA_001623345.3/ |
| *Physaria fendleri* | https://www.ncbi.nlm.nih.gov/datasets/genome/GCA_036630335.1/ |
| *Przewalskia tangutica* | https://www.ncbi.nlm.nih.gov/datasets/genome/GCA_029844215.1/ |
| *Pugionium cornutum* | https://www.ncbi.nlm.nih.gov/data-hub/genome/GCA_018901935.1/ |
| *Raphanus raphanistrum* subsp. *raphanistrum* | https://www.ncbi.nlm.nih.gov/datasets/genome/GCA_019706035.1/ |
| *Raphanus sativus* | https://www.ncbi.nlm.nih.gov/datasets/genome/GCF_000801105.2/ |
| *Rhododendron vialii* | https://www.ncbi.nlm.nih.gov/datasets/genome/GCA_030253575.1/ |
| *Schrenkiella parvula* | https://www.ncbi.nlm.nih.gov/datasets/genome/GCA_000218505.1/ |
| *Solanum dulcamara* | https://www.ncbi.nlm.nih.gov/data-hub/genome/GCA_947179165.1/ |
| *Solanum galapagense* | https://www.ncbi.nlm.nih.gov/data-hub/genome/GCA_025758845.1/ |
| *Solanum lycopersicoides* | https://www.ncbi.nlm.nih.gov/data-hub/genome/GCA_022817965.1/ |
| *Solanum lycopersicum* 5.0 | http://solomics.agis.org.cn/tomato/ftp/SL5.0_masked_genome/ |
| *Solanum melongena* | https://www.ncbi.nlm.nih.gov/data-hub/genome/GCA_000787875.1/ |
| *Solanum pennellii* | https://www.ncbi.nlm.nih.gov/data-hub/genome/GCF_001406875.1/ |
| *Solanum tuberosum* (Solyntus) | https://www.ncbi.nlm.nih.gov/data-hub/genome/GCA_015076265.1/ |
| *Swietenia macrophylla* | https://www.ncbi.nlm.nih.gov/datasets/genome/GCA_032401905.1/ |
| *Theobroma grandiflorum* | https://www.ncbi.nlm.nih.gov/datasets/genome/GCA_036013995.1/ |
| *Thlaspi arvense* | https://www.ncbi.nlm.nih.gov/data-hub/genome/GCA_911865555.2/ |
| *Arabidopsis thaliana* 1 | https://www.ncbi.nlm.nih.gov/data-hub/genome/GCF_0000001735.4/ |

**Supplementary Table 3.** Genomes for *S. tuberosum* alignment.

| Species | Link |
| --- | --- |
| *Anisodus acutangulus* | https://www.ncbi.nlm.nih.gov/datasets/genome/GCA_029783415.1/ |
| *Arabidopsis arenosa* | https://www.ncbi.nlm.nih.gov/datasets/genome/GCA_905216605.1/ |
| *Arabidopsis suecica* | https://www.ncbi.nlm.nih.gov/data-hub/genome/GCA_019202805.1/ |
| *Arabidopsis thaliana* 1 | https://www.ncbi.nlm.nih.gov/data-hub/genome/GCF_0000001735.4/ |
| *Arabis alpina* | https://www.ncbi.nlm.nih.gov/data-hub/genome/GCA_900128785.1/ |
| *Arabis montbretiana* | https://www.ncbi.nlm.nih.gov/data-hub/genome/GCA_001484125.2/ |
| *Brassica carinata* | https://www.ncbi.nlm.nih.gov/data-hub/genome/GCA_016771965.1/ |
| *Brassica juncea* | https://www.ncbi.nlm.nih.gov/data-hub/genome/GCA_018703725.1/ |
| *Brassica napus* | https://www.ncbi.nlm.nih.gov/data-hub/genome/GCF_020379485.1/ |
| *Brassica nigra* | https://www.ncbi.nlm.nih.gov/data-hub/genome/GCA_016432835.1/ |
| *Brassica oleracea* var. *oleracea* | https://www.ncbi.nlm.nih.gov/data-hub/genome/GCF_000695525.1/ |
| *Brassica rapa* | https://www.ncbi.nlm.nih.gov/data-hub/genome/GCF_000309985.2/ |
| *Camelina hispida* | https://www.ncbi.nlm.nih.gov/data-hub/genome/GCA_023657505.1/ |
| *Camelina laxa* | https://www.ncbi.nlm.nih.gov/data-hub/genome/GCA_024034495.1/ |
| *Camelina neglecta* | https://www.ncbi.nlm.nih.gov/data-hub/genome/GCA_023864065.1/ |
| *Camelina sativa* | https://www.ncbi.nlm.nih.gov/data-hub/genome/GCF_000633955.1/ |
| *Capsicum annum* | https://www.ncbi.nlm.nih.gov/data-hub/genome/GCF_002878395.1/ |
| *Capsicum baccatum* | https://www.ncbi.nlm.nih.gov/data-hub/genome/GCA_002271885.2/ |
| *Capsicum chinense* | https://www.ncbi.nlm.nih.gov/data-hub/genome/GCA_002271895.2/ |
| *Cucumis melo* | https://www.ncbi.nlm.nih.gov/datasets/genome/GCF_025177605.1/ |
| *Daucus carota* | https://www.ncbi.nlm.nih.gov/datasets/genome/GCA_030127425.1/ |
| *Erysimum cheiranthoides* | https://www.ncbi.nlm.nih.gov/data-hub/genome/GCA_011420285.1/ |
| *Euphorbia peplus* | https://www.ncbi.nlm.nih.gov/datasets/genome/GCA_028411795.1/ |
| *Gossypium herbaceum* | https://www.ncbi.nlm.nih.gov/datasets/genome/GCA_025502765.1/ |
| *Isatis tinctoria* | https://www.ncbi.nlm.nih.gov/data-hub/genome/GCA_010577795.1/ |
| *Lycium barbarum* | https://www.ncbi.nlm.nih.gov/data-hub/genome/GCA_019175385.1/ |
| *Lycium ferocissimum* | https://www.ncbi.nlm.nih.gov/datasets/genome/GCA_029784015.1/ |
| *Nicotiana attenuata* | https://www.ncbi.nlm.nih.gov/data-hub/genome/GCF_001879085.1/ |
| *Oryza sativa Indica Group* | https://www.ncbi.nlm.nih.gov/datasets/genome/GCA_001623345.3/ |
| *Przewalskia tangutica* | https://www.ncbi.nlm.nih.gov/datasets/genome/GCA_029844215.1/ |
| *Pugionium cornutum* | https://www.ncbi.nlm.nih.gov/data-hub/genome/GCA_018901935.1/ |
| *Rhododendron vialii* | https://www.ncbi.nlm.nih.gov/datasets/genome/GCA_030253575.1/ |
| *Solanum appendiculatum* | https://www.ncbi.nlm.nih.gov/data-hub/genome/GCA_018342035.1/ |
| *Solanum arcanum* | https://www.ncbi.nlm.nih.gov/datasets/genome/GCA_027789145.1/ |
| *Solanum chilense* | https://www.ncbi.nlm.nih.gov/data-hub/genome/GCA_006013705.1/ |
| *Solanum chmielewskii* | https://www.ncbi.nlm.nih.gov/datasets/genome/GCA_027704785.1/ |
| *Solanum clarkiae* | https://www.ncbi.nlm.nih.gov/data-hub/genome/GCA_011800125.1/ |
| *Solanum commersonii* | https://www.ncbi.nlm.nih.gov/data-hub/genome/GCA_018258275.1/ |
| *Solanum corneliomulleri* | https://www.ncbi.nlm.nih.gov/datasets/genome/GCA_027704805.1/ |
| *Solanum dulcamara* | https://www.ncbi.nlm.nih.gov/data-hub/genome/GCA_947179165.1/ |
| *Solanum galapagense* | https://www.ncbi.nlm.nih.gov/data-hub/genome/GCA_025758845.1/ |
| *Solanum habrochaites* | https://www.ncbi.nlm.nih.gov/datasets/genome/GCA_027704765.1/ |
| *Solanum lycopersicoides* | https://www.ncbi.nlm.nih.gov/data-hub/genome/GCA_022817965.1/ |
| *Solanum lycopersicum var. cerasiforme* | https://www.ncbi.nlm.nih.gov/datasets/genome/GCA_027705125.1/ |
| *Solanum lycopersicum var. M82* | https://www.ncbi.nlm.nih.gov/data-hub/genome/GCA_900008105.1/ |
| *Solanum melongena* | https://www.ncbi.nlm.nih.gov/data-hub/genome/GCA_000787875.1/ |
| *Solanum neorickii* | https://www.ncbi.nlm.nih.gov/datasets/genome/GCA_027704865.1/ |
| *Solanum okadae* | https://www.ncbi.nlm.nih.gov/data-hub/genome/GCA_023159065.1/ |
| *Solanum ossicruentum* | https://www.ncbi.nlm.nih.gov/data-hub/genome/GCA_025774065.1/ |
| *Solanum pennellii* | https://www.ncbi.nlm.nih.gov/data-hub/genome/GCF_001406875.1/ |
| *Solanum peruvianum* | https://www.ncbi.nlm.nih.gov/datasets/genome/GCA_027705065.1/ |
| *Solanum pimpinellifolium* | https://www.ncbi.nlm.nih.gov/data-hub/genome/GCA_014964335.1/ |
| *Solanum pinnatisectum* | https://www.ncbi.nlm.nih.gov/data-hub/genome/GCA_009887355.1/ |
| *Solanum rostratum* | https://www.ncbi.nlm.nih.gov/datasets/genome/GCA_029948455.1/ |
| *Solanum sejunctum* | https://www.ncbi.nlm.nih.gov/data-hub/genome/GCA_025642045.1/ |
| *Solanum sitiens* | https://www.ncbi.nlm.nih.gov/data-hub/genome/GCA_016801875.1/ |
| *Solanum stenotomum* | https://www.ncbi.nlm.nih.gov/data-hub/genome/GCF_019186545.1/ |
| *Solanum tuberosum* (DM 1-3 516 R44) | https://www.ncbi.nlm.nih.gov/data-hub/genome/GCF_000226075.1/ |
| *Solanum verrucosum* | https://www.ncbi.nlm.nih.gov/data-hub/genome/GCF_900185275.1/ |
| *Thlaspi arvense* | https://www.ncbi.nlm.nih.gov/data-hub/genome/GCA_911865555.2/ |

**Supplementary Table 4.** Intergenic region fractions for fixed-neutral and proxy-neutral sets for *thaliana* and *S. lycopersicum* datasets.

| **Species** | **Intergenic regions fraction**<br>**proxy-neutral** | **Intergenic regions fraction**<br>**fixed-neutral** |
| --- | --- | --- |
| *A. thaliana* | 0.272 | 0.271 |
| *S. lycopersicum* | 0.848 | 0.892 |

**Supplementary Table 5.**
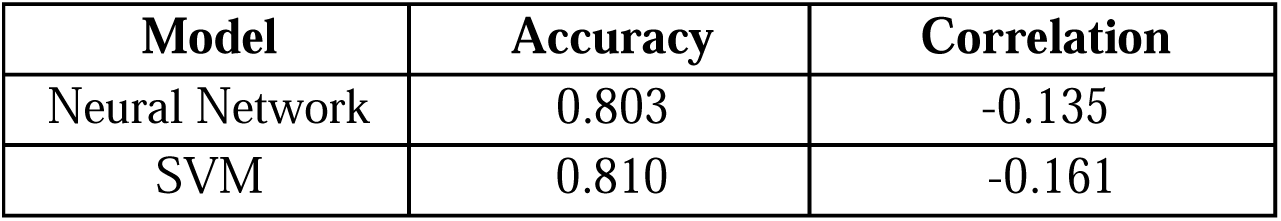
Metrics comparison between NN and SVM models for *S. lycopersicum*.

| **Model** | **Accuracy** | **Correlation** |
| --- | --- | --- |
| Neural Network | 0.803 | -0.135 |
| SVM | 0.810 | -0.161 |

**Supplementary Figure 1.**
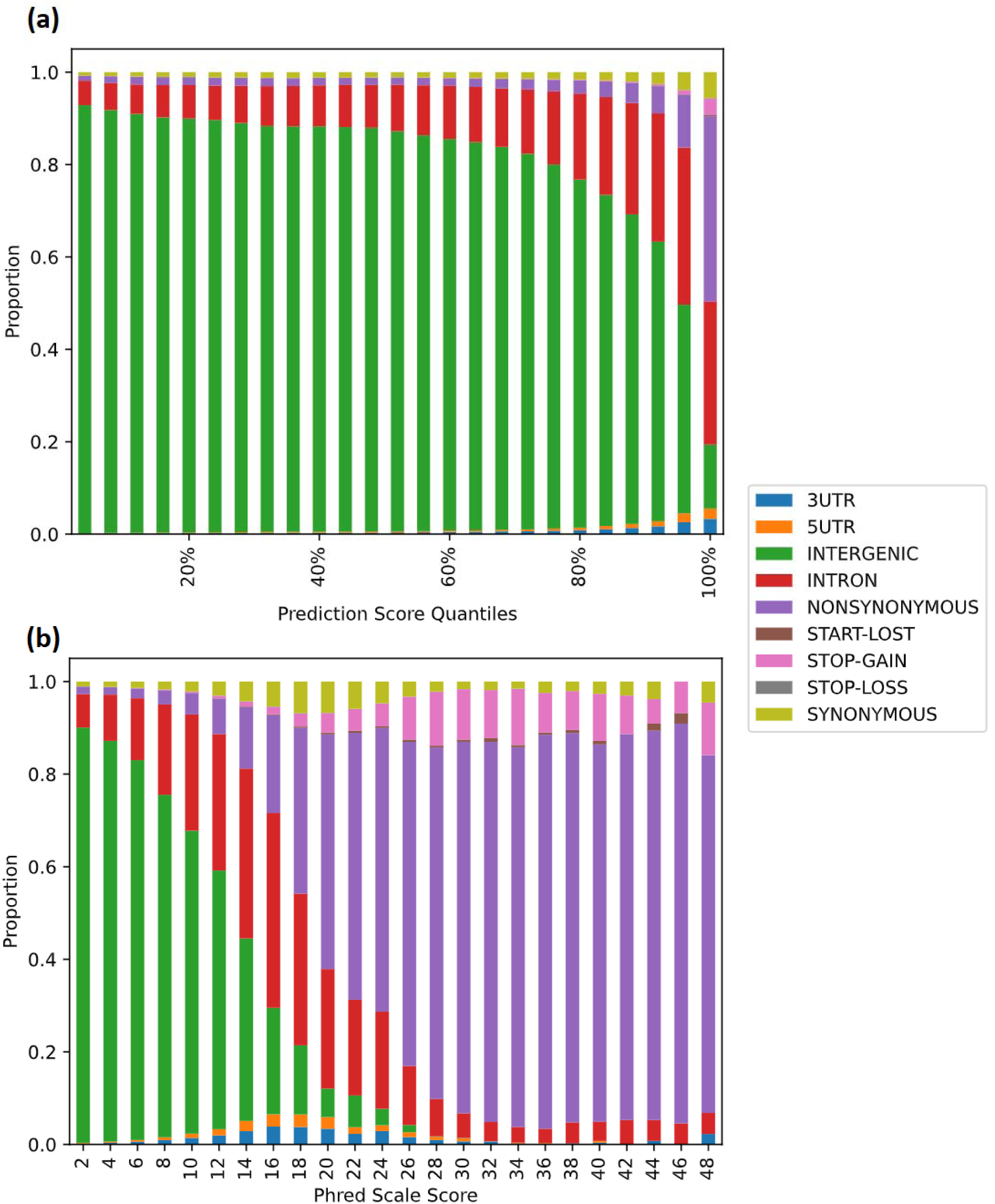
Validation plots for *S. lycopersicum* model trained with the fixed neutral set. Description is the same as Figure 3. The observed patterns appear very similar to what is observed in Figure 3 training on the whole neutral set.

**Supplementary Figure 2.**
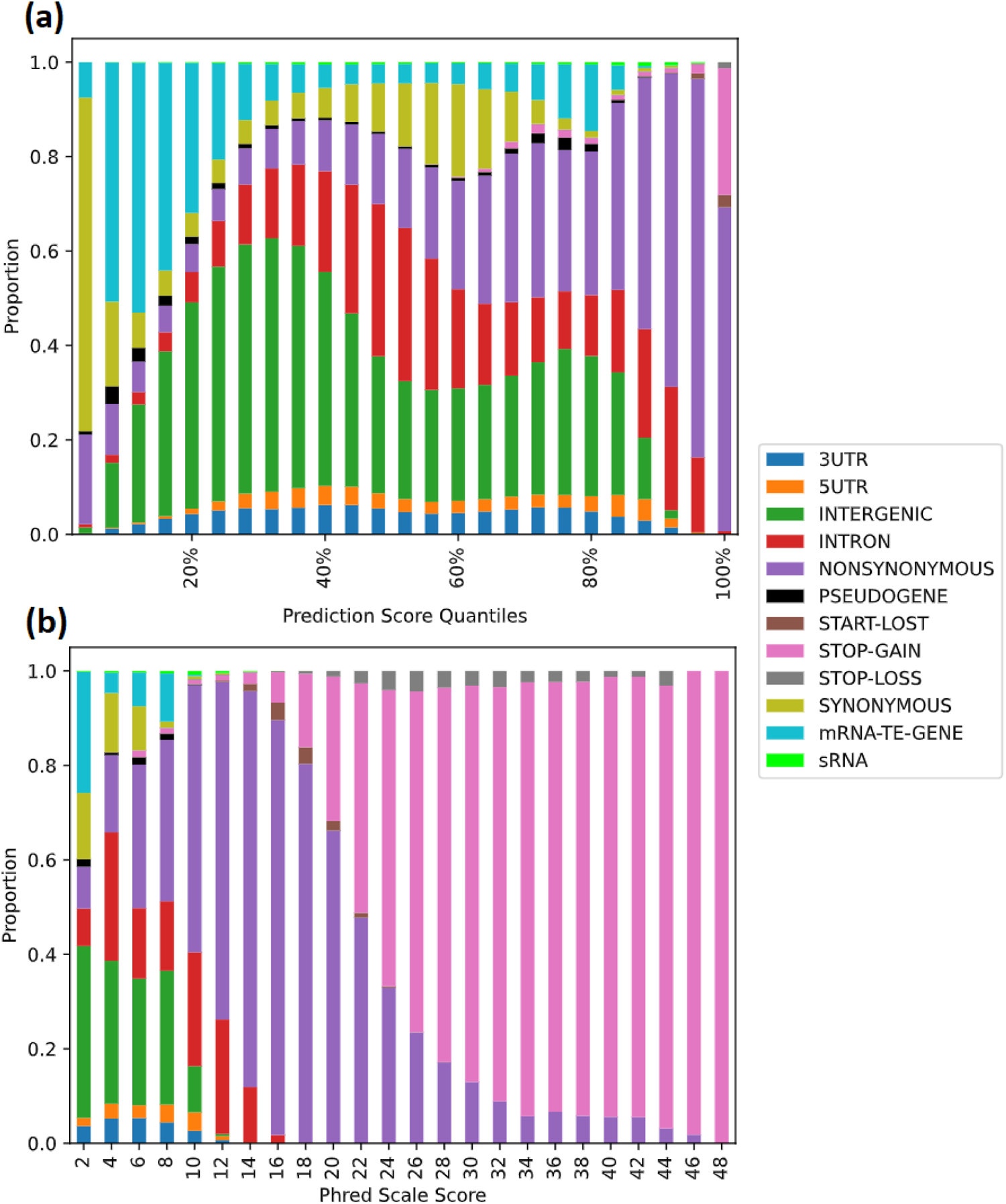
Validation plots for *A. thaliana* model trained with the fixed neutral set. Description is the same as Figure 4. The observed patterns appear very similar to what is observed in Figure 4 training on the whole neutral set.

